# Multiple motor proteins regulate ALS-linked TDP-43 anterograde axonal transport

**DOI:** 10.64898/2026.06.15.732214

**Authors:** Monica Feole, Victorio M. Pozo Devoto, Neda Dragišić, Maria Čarna, Katja Klosterman, Clara Limbaek-Stokin, Giancarlo Forte, Richard A. Smith, Clive N. Svendsen, Gorazd B. Stokin

## Abstract

TDP-43 is an RNA-binding protein essential for RNA metabolism. Under physiological conditions, it predominantly resides in the nucleus but is also expressed in the cytoplasm, where it regulates mRNA trafficking and local translation. In nearly 97% of Amyotrophic Lateral Sclerosis (ALS) cases, TDP-43 undergoes nuclear depletion and cytoplasmic aggregation. While its nuclear functions are well characterized, its axonal roles remain poorly understood, despite axonal degeneration being a hallmark of ALS pathology. To address this gap, we investigated TDP-43 localization and transport dynamics in axons of H9-derived human neurons. We compared fluorescently labeled TDP-43 with three well-characterized axonal cargoes, Rab5, synaptophysin, and APP, and examined protein-protein interactions between TDP-43 and the axonal transport machinery. Our analyses revealed that TDP-43 exhibits active anterograde axonal transport and interacts with multiple kinesin motor proteins, including all three KIF5 isoforms, through the adaptor KLC1, and the synaptic vesicles motor KIF1A. This multi-motor engagement suggests a flexible transport system that ensures mRNA delivery to distal axons. In ALS, where TDP-43 accumulates abnormally in the cytoplasm, this flexibility may become compromised, with multiple transport mechanisms simultaneously affected. This could contribute to progressive accumulation of non-functional TDP-43 granules, disrupting mRNA trafficking and local translation. Our findings provide a foundation for understanding how physiological TDP-43 transport mechanisms may be impaired during disease, highlighting axonal TDP-43 transport pathways as potential therapeutic targets.

## Introduction

TAR DNA-binding protein 43 (TDP-43) is a ubiquitously expressed and highly conserved RNA-binding protein (RBP) essential for RNA metabolism^1,2^. TDP-43 shuttles between the nucleus and cytoplasm, but predominantly localizes to the nucleus under physiological conditions^3,4^. Here, TDP-43 is an important regulator of transcription and splicing for a wide range of RNAs^5^. In the cytoplasm, TDP-43 supports trafficking and local translation of several mRNAs in proximal and distal axonal and dendritic compartments, including its own^6,7,8^.

TDP-43 pathology is a hallmark of several neurodegenerative diseases, most notably Amyotrophic Lateral Sclerosis (ALS) and Frontotemporal Lobar Degeneration (FTLD) and has also been observed in Alzheimer’s disease^9^, Huntington’s disease^10^, and others. In approximately 97% of sporadic ALS (sALS) cases, TDP-43 is depleted from the nucleus and abnormally accumulates in the cytoplasm, forming aggregates that can contribute to neuronal dysfunction and death^11,12,13^. This dual-pathology, characterized by nuclear depletion and cytoplasmic accumulation, disrupts TDP-43 function in both compartments. In the nucleus, TDP-43 normally regulates transcripts encoding STMN2 (stathmin-2), essential for axonal growth and microtubule dynamics, and UNC13A, which mediates synaptic vesicle priming at the neuromuscular junctions (NMJs)^14–16^. Loss of nuclear TDP-43 leads to cryptic exon splicing in these transcripts, resulting in non-functional proteins that compromise structural axonal stability and synaptic transmission^7,17–20^. Concurrently, cytoplasmic accumulation of TDP-43 impairs transport and local translation of mRNAs encoding cytoskeletal proteins as neurofilament light chain (NEFL)^21^, synaptic and nuclear-encoded mitochondrial mRNAs at neuromuscular junctions (NMJs)^6,22,23^, and ribosomal-associated protein mRNAs^24^, among others. These combined deficits, loss of nuclear splicing regulation and impaired cytoplasmic mRNA transport and translation, may drive the axonal degeneration characteristic of ALS.

In the cytoplasm, TDP-43 associates with specific mRNA pools to form ribonucleoprotein (RNP) complexes, which assemble into what are currently defined as membraneless organelles (MLOs)^25–28,29^. These MLOs undergo long-distance transport along microtubules, facilitated by motor and adaptor proteins^30,31^. Studies have revealed that some TDP-43-containing RNPs also ‘hitchhike’ on organelles to reach distal neuronal regions^32,33^. Though, whether TDP-43 containing RNPs rely solely on this non-canonical transport mechanism, *via* organelle interaction, or can directly interact with adaptor and motor proteins remains unclear.

Despite extensive characterization of TDP-43’s nuclear functions, its cytoplasmic role and axonal transport mechanisms remain poorly understood. It is unclear which motor proteins mediate TDP-43 trafficking in the axonal compartment, how this transport regulates mRNAs delivery to distal subdomains, and at what level the disruption of these mechanisms contributes to ALS pathology^34–36^. Understanding TDP-43 dynamics within axons may provide crucial insight into early pathogenic events and either support or challenge the *dying-back* model of ALS, in which distal axonal dysfunction and synaptic impairment precede and potentially drive motor neuron degeneration^37–39^.

To address these questions, we investigated the axonal transport dynamics of TDP-43 under physiological conditions. Using human neurons derived from the H9 embryonic stem cell line (H9-derived), we compared the movement of fluorescently labeled TDP-43 with three well-characterized axonal cargoes, Rab5, Synaptophysin (SYP), and Amyloid Precursor Protein (APP), each with distinct motor dependencies and directional bias linked to their role and localization in the axons. Through live-cell imaging, followed by segmental trajectory analysis, we demonstrated that TDP-43 exhibits processive anterograde transport with dynamics comparable to vesicular cargoes. Additionally, using both biochemical and proximity ligation approaches, we showed that TDP-43 engages with both anterograde and retrograde transport machinery, consistent with its bidirectional profile. Because anterograde transport is regulated by multiple kinesin isoforms^40^, while the retrograde transport is controlled by a single motor protein complex characterized by the dynein-dynactin complex^41,42^, we specifically examined which kinesin motors associate with TDP-43. We found that TDP-43 engages with multiple kinesin motor proteins, including all three kinesin-1 family isoforms, KIF5A, B, and C, through the adaptor KLC1, as well as the kinesin-3 family member KIF1A. In addition to previous finding describing alternative transport models^32,33,43^, this multi-motor engagement suggests a flexible TDP-43-containing MLOs transport system that ensures robust mRNA delivery to diverse axonal compartments, and that due to changes in TDP-43 fitness might be selectively affected under pathological conditions.

## Results

### Net TDP-43 axonal transport shows balanced proportions of anterograde and retrograde movement

While TDP-43 has been primarily associated with nuclear functions, it is also present in axons under physiological conditions where it regulates mRNA trafficking and local translation ^22,31,49,50^. To investigate TDP-43 axonal transport mechanisms, we performed live imaging experiments using previously characterized H9-derived human neurons for 40 days in vitro (DIV)^51^ **(Fig. 1A)**. Neurons were transfected with Lipofectamine 2000 mixed with plasmids expressing GFP-tagged cargoes: TDP-43, GRab5, Synaptophysin (SYP), and APP 48 hours prior to imaging **(Supplementary Fig. 1A)**. We compared TDP-43 with three established axonal cargoes (Rab5, SYP, and APP) that exhibit distinct transport properties ^40,52,53^. Movies were recorded at 4 frames per second for 60 seconds (s), and only axonal projections confirmed by post-imaging GFP/pNFH overlapping intensities were analyzed^44^ **(Supplementary Fig. 1B, Fig. 1B; Supplementary Movie 1, 2, 3, and 4)**.

**Figure 1.**
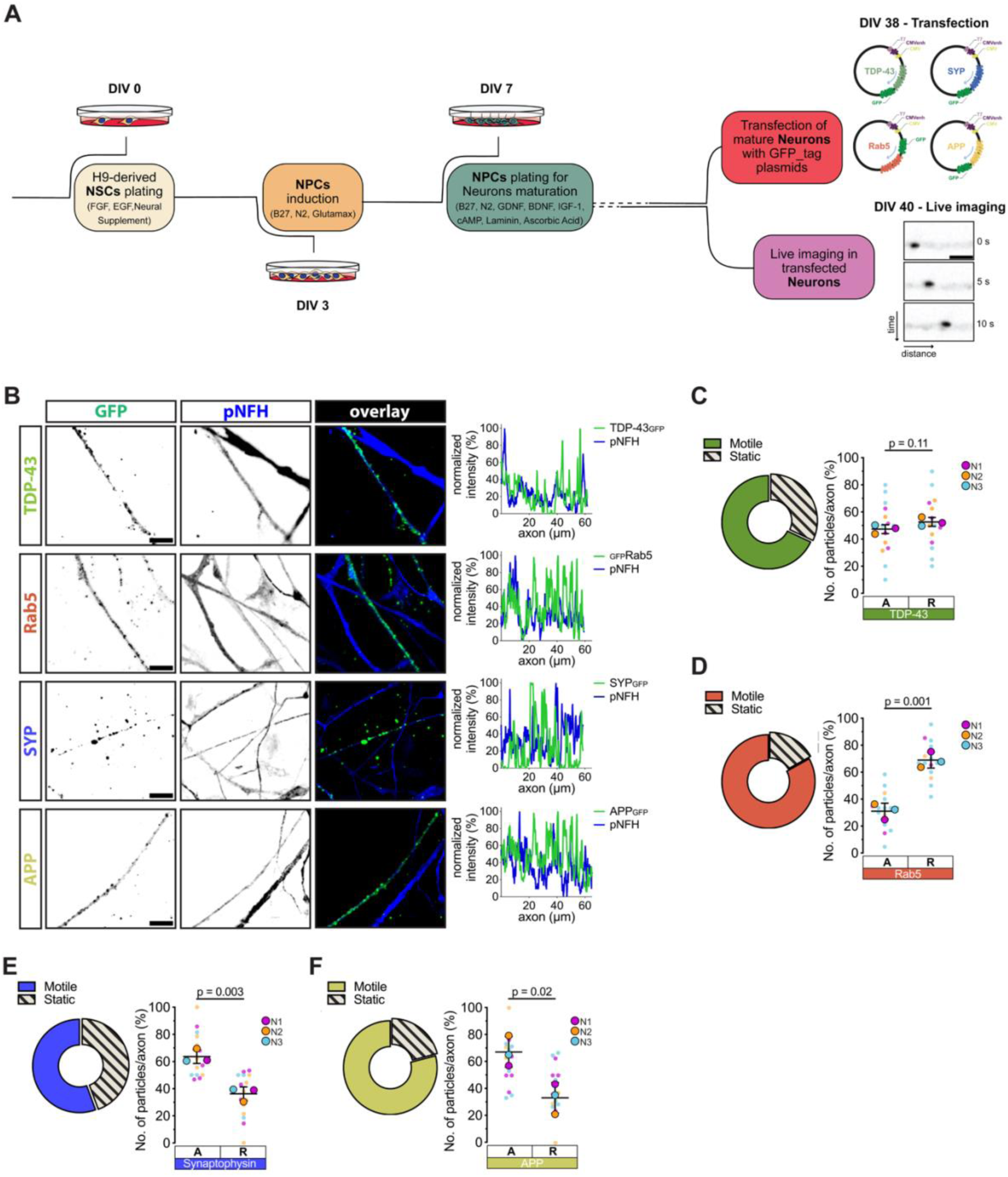
Axonal cargoes show distinct direction polarity. **(A)** Schematic representation of timeline differentiation of H9-derived neurons, and the two main time-points crucial for carrying axonal transport studies: DIV 38 – transfection of 4 different plasmids; DIV 40 recording of axonal cargoes movement. Micrographs in the bottom right represent screenshot of a particle moving over time. **(B)** Representative micrographs showing signal from GFP tag over-expressed cargoes, TDP-43, Rab5, SYP, and APP (green), with axonal pNFH marker (blue). On the right of the panel normalized intensities graphs show the overlapping signals in the illustrated ROIs (scale bar = 10µm). **(C - *left*)** Pie charts are showing the proportions of motile and completely static particles in the net axonal transport analysis of TDP-43; **(C - *right*)** superplots representing the proportions of motile TDP-43 particles moving anterograde and retrograde (*n*=15 axons from *n*=3 biological replicates). **(D - *left*)** Pie charts are showing the proportions of motile and completely static particles in the net axonal transport analysis of Rab5; **(D - *right*)** superplots representing the proportions of motile Rab5 particles moving anterograde and retrograde (*n*=15 axons from *n*=3 biological replicates). **(E - *left*)** Pie charts are showing the proportions of motile and completely static particles in the net axonal transport analysis of SYP; **(E - *right*)** superplots representing the proportions of motile SYP particles moving anterograde and retrograde (*n*=15 axons from *n*=3 biological replicates). **(F - *left*)** Pie charts are showing the proportions of motile and completely static particles in the net axonal transport analysis of APP; **(F - *right*)** superplots representing the proportions of motile APP particles moving anterograde and retrograde (*n*=15 axons from *n*=3 biological replicates). Data show normalized intensities along the GFP(+) and pNFH(+) ROIs **(B)**, pie charts from *n*=3 biological replicate means per each cargo **(C, D, E, and F – *left*)**, mean ± S.D. **(C, D, E, and F – *right*)**. Statistical analyses were performed using unpaired *t*-test, and comparisons result shown with exact *p* values.

Using semi-automated spot-tracking, we first confirmed that all four cargoes showed comparable GFP signal levels and track densities, validating the reliability of subsequent transport comparisons only in axonal projections **(Supplementary Table 2, Fig. 1B - *right*, Supplementary Fig.1C)**. We then assessed the proportion of motile versus static particles using a uniform movement threshold applied across all cargoes to enable direct comparison. This revealed that 50-80% of particles were motile while 20-50% remained static for all four proteins **(*pie charts* – Fig. 1C, 1D, 1E, and 1F)**. Because each cargo has distinct cargo-specific transport dynamics, this uniform threshold necessarily captures particles with varying velocities, including slower-moving fractions and particles undergoing pausing or directional changes within the motile category.

Next, we assessed within the motile fractions the proportions of particles that for each cargo were moving toward the synaptic terminals (anterograde) or toward the soma (retrograde). TDP-43 exhibited balanced bidirectional transport with no significant directional preference **(Fig. 1C - *right*)**, suggesting widespread axonal distribution similar to other mRNA-trafficking RBPs^21,22,54^. In contrast, the other cargoes showed their canonical transport patterns: Rab5 displayed predominantly retrograde movement consistent with its endosomal recycling function^55,56^ **(Fig. 1D - *right*)**. SYP showed strong anterograde bias reflecting kinesin-3/KIF1A-mediated transport to synapses^57,58^ (**Fig. 1E - *right*)**. APP exhibited slightly anterograde-biased but relatively balanced bidirectional movement^44,59–61^ **(Fig. 1F - *right*)**. These distinct transport dynamics support the involvement of multiple motor proteins and adaptor complexes in cargo-specific axonal transport, which in the case of TDP-43 seems to be equally important for its wide distribution along the axon.

### TDP-43 exhibits fast axonal transport comparable to vesicle-bound cargoes SYP and APP

To characterize the transport mechanisms of individual cargo trajectories, we employed a ’segmental analysis’ approach that divides each trajectory into vectorial segments, allowing detailed examination of movement properties including direction, velocity, and pausing behavior by collecting real-time data from the Imaris software^44,51^ **(Fig. 2A – *top*)**. This frame-by-frame analysis reveals which motor proteins or adaptor complexes may mediate microtubule-directed transport of specific cargoes while also enabling categorization of trajectory types^47,62^ **(Fig. 2A – *bottom*)**. To examine the regulatory aspects of distinct motor protein-cargo interactions, we chose to include only motile particles in the segmental analysis (**Supplementary Table 3)**.

**Figure 2.**
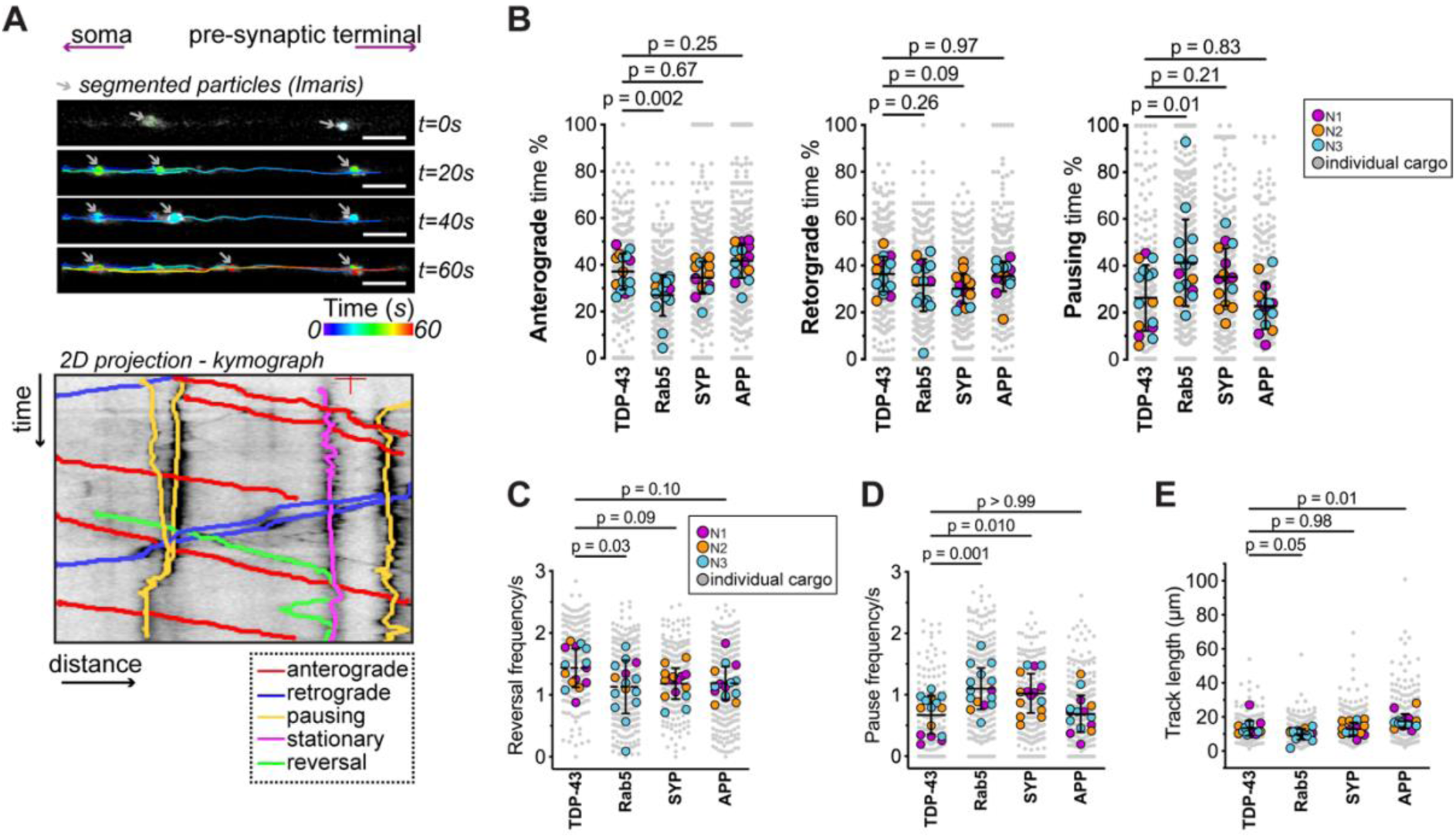
TDP-43 exhibits active anterograde axonal transport similar to SYP and APP. (A - *top*) Representative micrographs of particles movement, segmentation and tracking determined with Imaris spots tracking algorithms. Grey arrows follow the particle movement over-time, as indicated by the time points illustrated on the right of the micrographs, while colored time scale bar indicates the length of movement as illustrated by the colored trajectories in the micrographs (scale bar = 5µm); **(A - *bottom*)** representative 2D projection (kymograph) of possible particles behavior (y=time, x=distance); trajectories can be traced to describe either motion towards the synaptic terminal (anterograde - red), to the soma (retrograde), no clear direction bias (pausing cargo – yellow), not moving cargo (stationary – magenta), or changing directionality (reversing cargo – green). **(B)** Quantification and comparisons of real time movement analyzed as proportion of time spent by each TDP-43, Rab5, SYP, and APP particle in anterograde (left), retrograde (center), or pausing (right). Superplots represent single trajectories of particles (opaque colored dots) superimposed by means per axon (solid-colored circles), with colors and legends depicting the biological replicate to which every dot belongs (*n*>360 tracks from *n*=15 axons per each cargo analyzed from 3 different biological replicates). **(C)** Quantification and comparisons of reversal frequency normalized per second of motile TDP-43, Rab5, SYP, and APP particles (*n*>250 tracks from *n*=15 axons per each cargo analyzed from 3 different biological replicates). **(D)** Quantification and comparisons of pause frequency normalized per second of motile TDP-43, Rab5, SYP, and APP particles (*n*>250 tracks from *n*=15 axons per each cargo analyzed from 3 different biological replicates). **(E)** Quantification and comparisons of total track lengths of motile TDP-43, Rab5, SYP, and APP particles moving either anterograde or retrograde (*n*>250 tracks from *n*=15 axons per each cargo analyzed from 3 different biological replicates). Data show mean ± S.D. of superplot with axon means over the single tracks’ dots **(B, C, D, and E).** Statistical analyses were performed using one-way ANOVA followed by Dunnet’s multiple comparisons test **(B)**, or Kruskal Wallis test followed by Dunn’s post hoc test **(C, D, and E)**, and the corresponding results shown with exact *p* values.

Segmental directionality analysis revealed that TDP-43 spent a significantly greater proportion of time in anterograde movement compared to Rab5 but showed similar anterograde proportions to SYP and APP **(Fig. 2B - *left*)**. For retrograde movement, TDP-43 was comparable to Rab5, SYP and APP **(Fig. 2B - *center*)**. Pausing time was significantly lower in TDP-43 compared to Rab5, with no differences observed relative to SYP or APP **(Fig. 2B - *right*)**. These real-time motion patterns indicate that TDP-43 shares transport dynamics with the fast-moving cargoes SYP and APP rather than the retrograde-biased Rab5.

We next quantified directional reversals (frequency of direction changes during transport) and found that TDP-43 reversed direction significantly more frequently than Rab5, while showing similar reversal rates to SYP and APP **(Fig. 2C)**. Similarly, pause frequency (number of pausing events per trajectory) was significantly lower for TDP-43 compared to Rab5 and SYP, but comparable to APP **(Fig. 2D)**. Finally, we measured total track length as an indicator of cargo processivity, which reflects the ability of a motor protein to step along microtubules in a constant way before detaching from the cytoskeletal track. TDP-43 particles traveled distances comparable to SYP and Rab5, and exhibited significantly shorter track lengths compared to APP, which displayed the widest range of distances **(Fig. 2E)**. This suggests that TDP-43 transport may be characterized by both short- and long-distance movements, reflecting potential heterogeneity in cargo composition or motor protein engagement.

Collectively, these segmental analyses demonstrate that TDP-43-containing MLOs, can be considered as fast axonal cargoes with transport characteristics resembling the vesicle-associated proteins SYP and APP. This is particularly noteworthy because classical models of axonal transport have been predominantly developed from studies of membrane-bound vesicular cargoes, where transmembrane proteins serve as adaptor binding sites for motor protein recruitment and cargo sorting at axonal entry points^63,64^.

### TDP-43 anterograde transport combines SYP-like processivity with APP-matching velocities

To determine whether longer uninterrupted transport events correlate with sustained velocity increase, we analyzed segmental velocities for all four cargoes during both anterograde and retrograde movement. First, relationship between segment duration and velocity was determined, which would indicate processive motor engagement. Correlation analysis using cubic spline fitting revealed cargo-specific patterns both in anterograde and retrograde direction.

For both TDP-43 and Rab5 segmental velocities correlated weakly but significantly with segment duration for both anterograde and retrograde transport **(Fig. 3A, and 3B)**. The correlation strengths did not differ significantly between directions, indicating that TDP-43 cargoes exhibit similar coupling of run length and speed in both directions **(Supplementary Fig. 2A)**.

**Figure 3.**
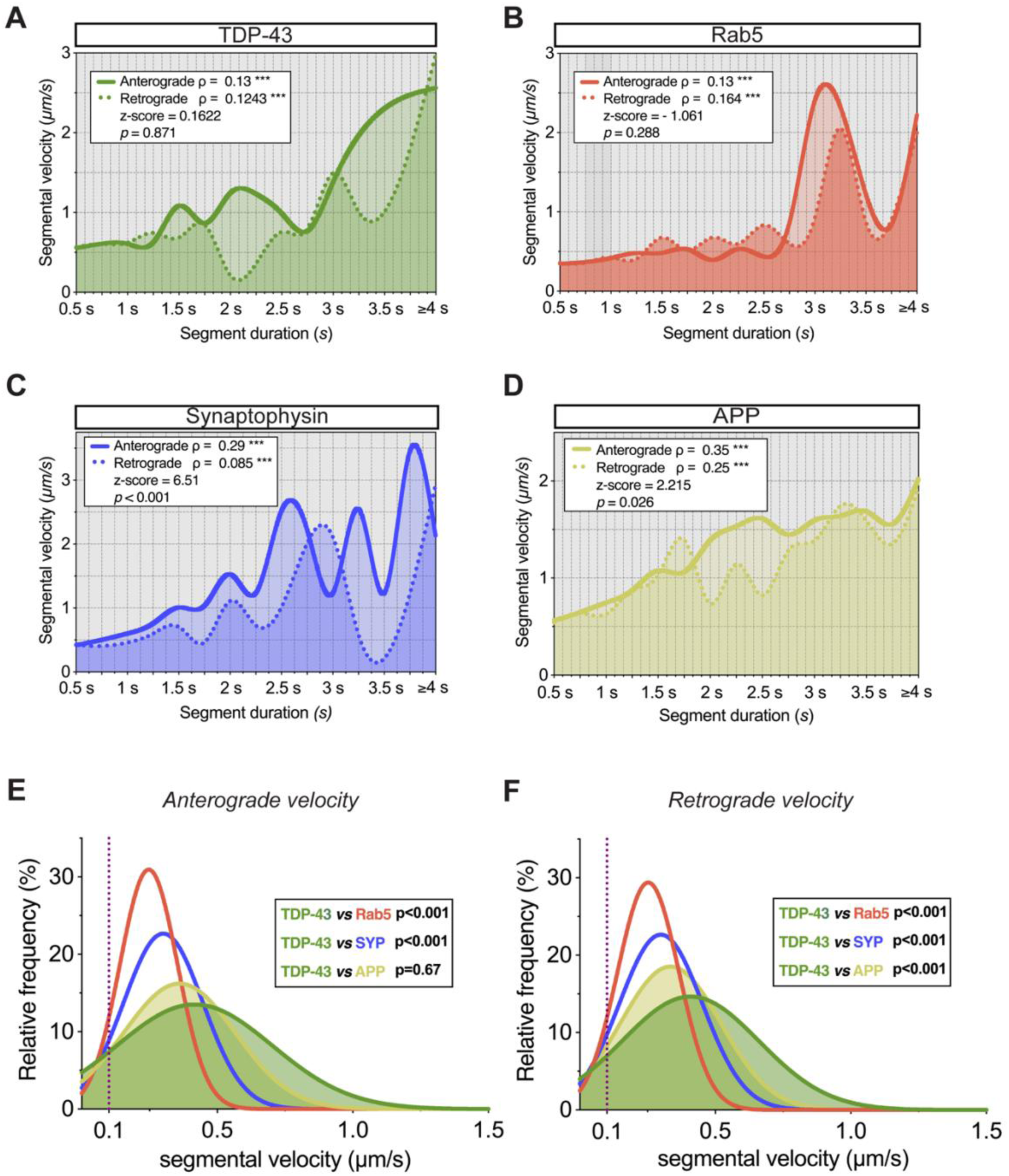
Positive correlations between velocity and run lengths represent processive anterograde and retrograde TDP-43 movement. Cubic spline fitting of correlation analyses for both anterograde and retrograde segmental velocities *vs* run length (segment duration) for TDP-43 **(A)**, Rab5 **(B)**, SYP **(C)**, and APP **(D)**. Graph shows fitting of all analyzed segments for anterograde (solid line) and retrograde (dashed line) moving TDP-43 particles, with Spearman’s correlation (𝛒) values and z-scores testing the correlation strength in either anterograde or retrograde (*n* >1500 segments analyzed from 15 axons from 3 biological replicates). **(E)** Frequency distribution of anterograde segmental velocity compared across the 4 cargoes TDP-43, Rab5, SYP, and APP, with colored AUC of APP and TDP-43 frequencies representing the overlap between TDP-43 and APP velocities (*n* >1500 segments analyzed from 15 axons from 3 biological replicates). **(F)** Frequency distribution of retrograde segmental velocity compared across the 4 cargoes TDP-43, Rab5, SYP, and APP, with colored AUC of APP and TDP-43 frequencies representing the overlap between TDP-43 and APP velocities (*n* >1500 segments analyzed from 15 axons from 3 biological replicates). Data are shown as cubic spline fitting models, where each point is crossed by fitting line curve **(A, B, C, and D)**, and frequency distributions in percentage of segmental velocity values **(E, and F)**. Statistical analyses were performed by using Spearman’s correlation and Fisher’s z-transformed then compared with a two-tailed z-test **(A-D)** Spearman’s 𝛒 values and z-scores are stated in the inlet tables, as well as corresponding significance (\*\*\**p* <0.001). Kruskal-Wallis followed by Dunn’s multiple comparisons test was used for comparison between frequency distributions **(E, and F)**.

In the case of SYP, segmental velocity correlated positively with segment duration for both anterograde- and retrograde-biased transport, but the anterograde correlation was significantly stronger, indicating direction-specific coupling between segment duration and velocity for SYP-positive vesicles **(Fig. 3C, and Supplementary Fig. 2A)**. Finally, we examined APP segmental velocities and observed that they also correlated positively with segment duration for both anterograde and retrograde runs which was significantly stronger for anterograde transport, indicating that APP cargoes exhibit a higher degree of coupling between run length and velocity during transport toward pre-synaptic terminals.

We then compared analysis of velocity distributions, which provided insight into potential motor protein involvement. For anterograde movement, TDP-43 velocities were significantly higher than those of Rab5 and SYP, while they showed near-complete overlap with APP anterograde velocities **(Fig. 3E, Supplementary Fig. 2B)**. This striking similarity between TDP-43 and APP anterograde velocity distributions suggests they may share common motor complexes or regulatory mechanisms for plus-end-directed transport. In contrast, retrograde segmental velocities of TDP-43 were significantly different from all three reference cargoes **(Fig. 3F, Supplementary Fig. 2C)**. This suggests that while the dynein-dynactin complex mediates retrograde transport for all cargoes, it exhibits cargo-specific processivity regulation^42,63,65^. The distinct retrograde velocity profile of TDP-43 MLOs may reflect unique adaptor proteins or post-translational modifications that modulate dynein-dynactin activity on membraneless RNP granules compared to vesicular cargoes^66^.

Collectively, these velocity analyses support the hypothesis that TDP-43 anterograde transport is mediated by kinesin family members similar to those driving APP and SYP movement. The striking similarities in TDP-43 anterograde transport characteristics, including processivity and velocity profiles, suggest that TDP-43-containing MLOs may utilize overlapping motor mechanisms to reach distal axonal compartments and synaptic regions. This would be functionally significant, as TDP-43-mediated local mRNA translation at synapses is crucial for maintaining synaptic plasticity and neuronal integrity^18,22,23,67^.

### TDP-43 engages with KLC1 and DCTN1 adaptors for effective anterograde and retrograde transport

To identify the molecular machinery underlying TDP-43 axonal transport, we performed co-immunoprecipitation (co-IP) experiments in 40 DIV human neurons, using total protein lysates cleaned from the nuclear fraction. Our experiments focused on kinesin light chain 1 (KLC1), an adaptor for KIF5-mediated anterograde transport ^68,69^, and dynactin 1 (DCTN1), a key component of the retrograde transport dynein complex ^70,71^ **(Fig. 4A)**.

**Figure 4.**
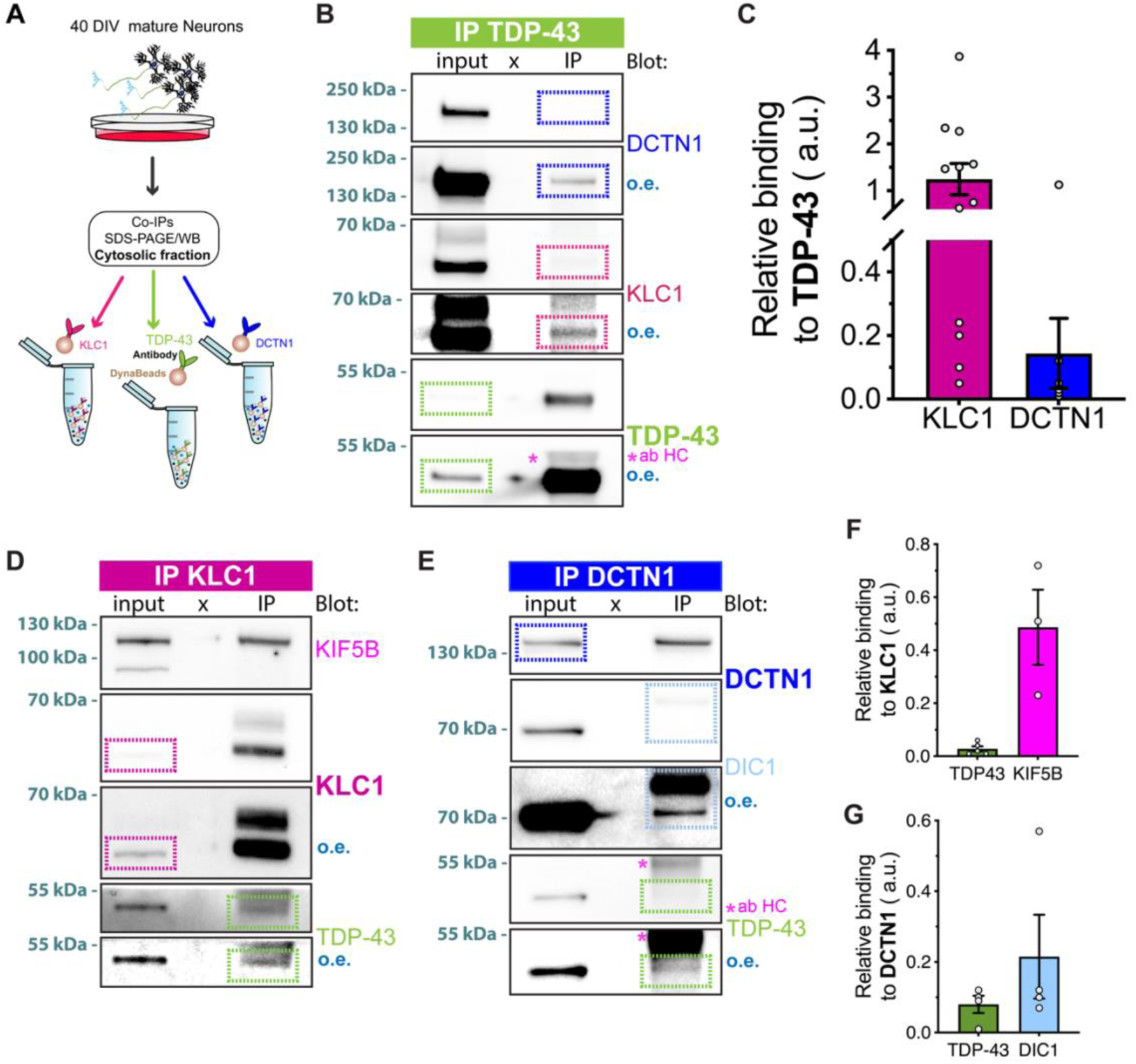
TDP-43 axonal transport is sustained by interaction with anterograde KLC1 and retrograde DCTN1 motor protein adaptors. **(A)** Schematic representation of co-IP experiments performed in 40 DIV human neurons to validate interactions between TDP-43, KLC1, and DCTN1. **(B)** Representative immunoblots of TDP-43 IP bands (green box) and DCTN1(blue box) and KLC1 (dark magenta box) co-IP bands. Each protein is represented by two images to show both unsaturated and over-exposed images (o.e.). **(C)** Co-IPs ratios expressed as KLC1 and DCTN1 intensity relative to TDP-43 IP (IP TDP-43 *n*=12 biological replicates with KLC1 co-IP analyzed in *n*=12 and DCTN co-IP intensities analyzed in *n*=10 biological replicates). **(D)** Representative immunoblots of KLC1 IP bands (dark magenta box) and KIF5B (magenta box) and TDP-43 (green box) co-IP bands. Each protein is represented by two images to show both unsaturated and over-exposed images (o.e.). **(E)** Co-IPs ratios expressed as TDP-43 and KIF5B intensity relative to KCL1 IP (IP KCL1 *n*=5 biological replicates with TDP-43 co-IP analyzed in *n*=5 and KIF5B co-IP intensities analyzed in *n*=3 biological replicates). **(F)** Representative immunoblots of DCTN1 IP bands (blue box) and DIC1 (light blue box) and TDP-43 (green box) co-IP bands. Each protein is represented by two images to show both unsaturated and over-exposed images (o.e.). **(G)** Co-IPs ratios expressed as TDP-43 and DIC1 intensity relative to DCTN1 IP (*n*=5 biological replicates).

IP of endogenous TDP-43 revealed positive interactions with both KLC1 and DCTN1 **(Fig. 4B, Supplementary Fig. 3A)**. Quantification of relative binding ratios to TDP-43 exhibited a more robust association with KLC1 rather than DCTN1 **(Fig. 4C)**. This preferential binding to the anterograde adaptor is consistent with our live-imaging data showing that TDP-43 exhibits fast anterograde transport characteristics and spends considerable time moving toward distal axonal regions.

To confirm the specificity of these interactions, we performed reciprocal IPs. Pulldown of KLC1 successfully co-IPed TDP-43, along with the expected positive control KIF5B, which forms the canonical KLC1-KIF5 motor complex **(Fig. 4D, Supplementary Fig. 3B)**. Although KLC1 is an abundant neuronal protein involved in transporting multiple cargoes, quantification indicated that a fraction of TDP-43 associates with KLC1, suggesting this interaction is physiologically relevant **(Fig. 4F)**. Similarly, IP of DCTN1 co-purified TDP-43 along with dynein intermediate chain 1 (DIC1), a core component of the dynein-dynactin complex^48,72^ **(Fig. 4E, Supplementary Fig. 3C)**. Binding ratio analysis confirmed that TDP-43 associates with DCTN1-containing complexes, supporting dynein-dynactin involvement in retrograde TDP-43 transport **(Fig. 4G)**.

Collectively, these biochemical interactions, in addition to the axonal transport data, demonstrate that TDP-43 engages with both anterograde (KLC1/KIF5) and retrograde (DCTN1/dynein) motor complexes, with a strong association with the anterograde machinery. The presence of both interactions provides a molecular explanation for the bidirectional transport dynamics observed in our live-imaging analyses and implicates KIF5 as an important kinesin mediating TDP-43 anterograde movement.

### TDP-43 associates with KLC1 and all three KIF5 isoforms in the axonal compartment

To validate the TDP-43-KLC1 interaction *in situ* and assess its functional specificity, we performed proximity ligation assays (PLAs) in our 40 DIV human neurons. PLA detects protein-protein interactions within a distance of ≤ 40 nm, allowing visualization of endogenous complexes in their native axonal environment.

We first established assay specificity using biological controls. For TDP-43, we used UBQLN2 as a positive control (known to bind TDP-43’s C-terminus^73^) and hnRNPC as a negative control (binds TDP-43 predominantly in the nucleus^74^) **(Supplementary Fig. 4A)**. For KLC1, we used kinesin heavy chain (KHC/KIF5) as a positive control^54^ and MyoV (an actin-based motor^40^) as a negative control, which showed expected positive and negative overlapping intensity patterns, respectively **(Supplementary Fig. 4B)**. We then validated the overlapping signals for TDP-43 and KLC1 antibodies with the axonal marker pNFH to confirm their suitability for PLA **(Supplementary Fig. 4C)**. Moreover, the KLC1 antibody alone showed no background binding in pilot PLA experiments and no overlap with pNFH **(Supplementary Fig. 4D)**. Having validated assay performance and antibody specificity, we next examined the TDP-43-KLC1 interaction directly.

To test TDP-43-KLC1 binding we performed shRNA-mediated KLC1 knockdown from day 16 to day 40 *in vitro* human neurons, comparing three conditions: non-transduced (NT), scramble_tGFP (scramble), and shRNA_tGFP KLC1 (shRNA KLC1) **(Fig, 5A, Supplementary Fig. 5E)**. KLC1 downregulation was induced with shRNAs of different efficiency, validated by western blot and densitometric analysis. First, we confirmed that βIII-tubulin loading control levels were comparable across all samples, with no significant differences found, suggesting that the treatment was not affecting protein levels other than the KLC1 **(Supplementary Fig. 5F)**. Next, we compared the levels of transduced scramble or different shRNA KLC1 constructs. By performing densitometric analysis and normalizing KLC1 levels to βIII-tubulin and tGFP, we observed that the most significant reduction was obtained with shRNA KLC1 A, which was selected for PLA analysis **(Supplementary Fig. 5G)**. We then performed PLAs, analyzing all conditions in pNFH-(+) axonal compartments. Neurons non-transduced (NT) and treated with control constructs displayed robust PLA signals along axons, confirming TDP-43-KLC1 association in the axonal compartment (**Fig. 5B - *left, center***). In contrast, shRNA KLC1-treated neurons had dramatically reduced PLA signals in tGFP-(+) axons (**Fig. 5B - *right***). Quantification revealed significantly higher PLA puncta density in both NT and scramble neurons compared to shRNA KLC1-treated neurons, with no significant difference between NT and scramble conditions **(Fig. 5C)**. These results demonstrate that reduced KLC1 levels specifically decrease TDP-43-KLC1 interaction, validating the specificity of this association.

**Figure 5.**
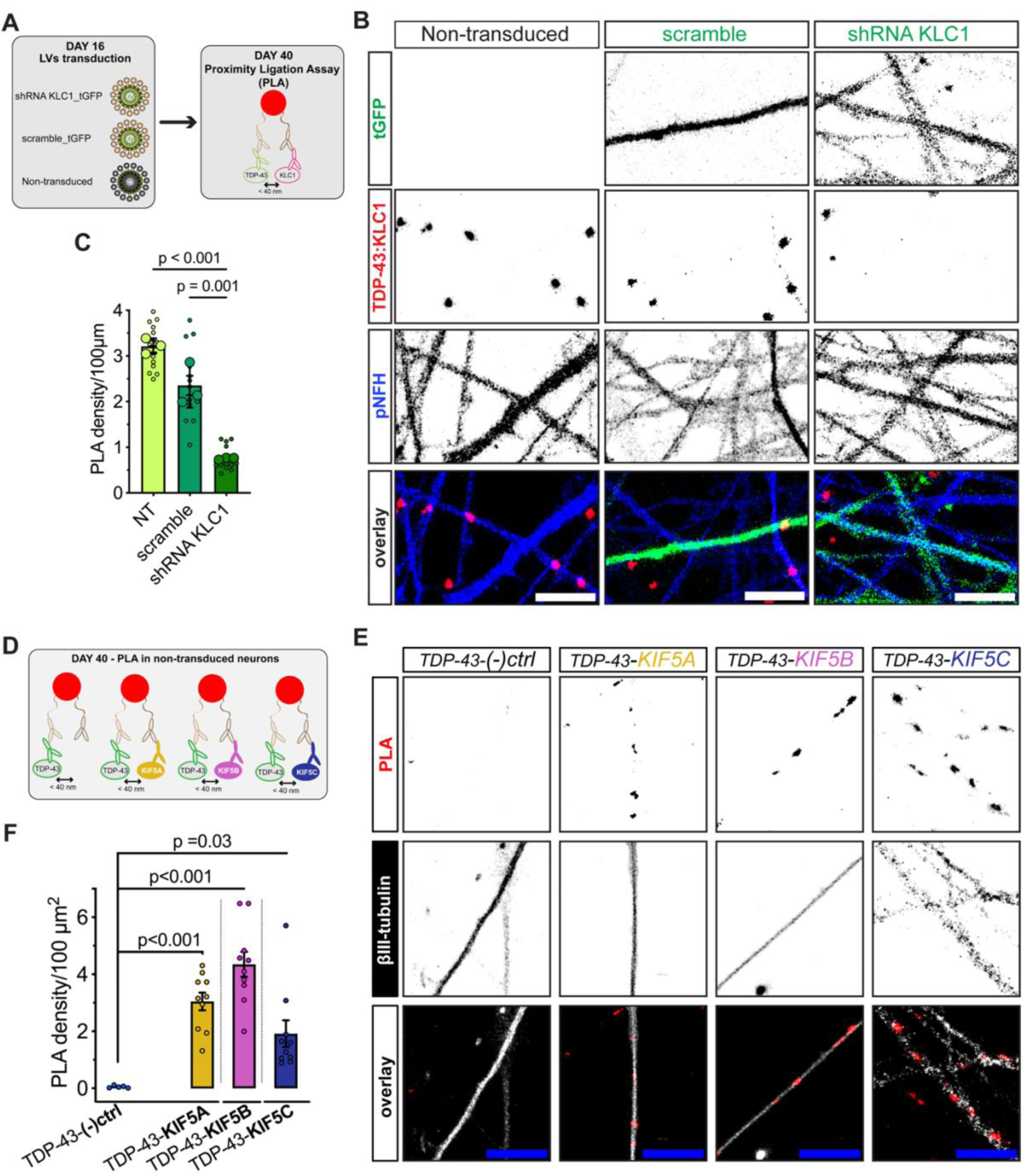
KLC1 and the three KIF5 isoforms associate with TDP-43 sustaining anterograde axonal transport. **(A)** Schematic representation of LV transduction strategy for KLC1 downregulation experiments, followed by PLA at DIV 40. **(B)** Representative micrographs of non-transduced, scramble GFP transduced, and shRNA KLC1 transduced neurons showing GFP signals (green), in scramble and shRNA, PLA dots representing positive/negative interaction between TDP-43:KLC1 (red), axonal marker pNFH (blue) and overlay images (scale bar = 10 µm). **(C)** Quantification of PLA densities analyzed along axonal projections (pNFH) performed in NT, scramble, and shRNA KLC1 neurons (*n*=14 images for NT and scramble and *n*=16 images for shRNA KLC1 from 3 different biological replicates; mean values per image were obtained from 7≤n≤24 traced axons per image). **(D)** Schematic representation of PLA experiments conducted at DIV 40 in human neurons using specific KIF5A, B, and C antibodies to detect interaction with TDP-43. **(E)** Representative micrographs of the PLA experiments performed to validate the interaction of TDP-43 with the three different isoforms of motor protein family kinesin-1 including KIF5A, KIF5B, and KIF5C. Images show (+) PLA signal (red), and the axonal marker βIII-tubulin (white), along which is possible to follow the red spots. Pictures show negative control with TDP-43 antibody only (left), TDP-43-KIF5A (center left), TDP-43-KIF5B (center right), and TDP-43-KIF5C (right); last row represents overlay images of PLA signal along neuronal projections (scale bar = 10 µm). **(F)** Quantification of PLA densities in TDP-43:KIF5A, TDP-43:KIF5B, and TDP-43:KIF5C *vs* negative control (*n*=5 images and *n*=10 images from 3 different biological replicates for TDP-43-(-)ctrl and TDP-43:KIF5A/B/C PLAs, respectively). Data show density of PLA dots over 100 µm axonal distance as mean ± S.D (**C)**, and density of PLA dots over 100 µm^2^ as mean ± S.D **(F)**. Statistical comparison was performed using a One-way ANOVA followed by Tukey’s multiple comparisons test **(C**) and unpaired *t*-test with Welch’s correction for independent comparisons since different antibodies cannot be compared between each other, though TDP-43-(-)ctrl was run in parallel **(F)**. Approximate or exact *p* values are shown in the figure.

Having confirmed TDP-43 interacts with KLC1, we next investigated which kinesin-1 heavy chain isoform associates with TDP-43. The mammalian kinesin-1 family includes three isoforms, KIF5A, KIF5B, and KIF5C, that are differentially expressed in different tissues and can exhibit cargo specificity^75^. KIF5B is ubiquitously expressed, while KIF5A and KIF5C are neuron-specific, making all three relevant candidates for neuronal TDP-43 transport. We performed PLA in 40 DIV neurons using isoform-specific antibodies against each KIF5 variant and TDP-43, with a βIII-tubulin_GFP cytoskeletal tracker marking neurite compartments **(Fig.e 5D, Supplementary Fig. 5A)**. Remarkably, confocal imaging revealed positive PLA puncta for all three isoforms with TDP-43 **(Fig. 5E)**, and quantification confirmed significantly elevated PLA puncta for TDP-43 with KIF5A, KIF5B, and KIF5C compared to negative controls **(Fig. 5F)**. This suggests that TDP-43 granules can engage with multiple kinesin-1 isoforms, which may contribute to the robust and heterogeneous anterograde transport observed in our live-imaging analyses.

Collectively, these PLA experiments, combined with KLC1 knockdown, validate the biochemical interaction data and demonstrate that TDP-43 specifically associates with the KLC1/KIF5 anterograde transport machinery in axons. The engagement with all three KIF5 isoforms is consistent with TDP-43’s robust anterograde transport characteristics. However, whether TDP-43 exhibits preferential binding to specific isoforms for delivery of specific mRNAs to distinct axonal subdomains remains to be determined.

### TDP-43 is alternatively recruited by the synaptic vesicle motor KIF1A

Our axonal transport analyses revealed that TDP-43 shares several dynamic characteristics with SYP, including processive anterograde runs and comparable overall transport behavior. SYP is primarily transported by KIF1A, a kinesin-3 family member specialized for fast anterograde delivery of synaptic vesicle precursors to presynaptic terminals^76,77^. Given these transport similarities, we hypothesized that KIF1A might also contribute to TDP-43 anterograde trafficking, potentially providing an additional or complementary transport pathway to the KIF5/KLC1 mechanism.

To test this hypothesis, we performed co-IPs and PLAs in 40-DIV human neurons **(Figure 6A)**. IP of TDP-43 successfully co-purified KIF1A **(Fig. 6B, and Supplementary Fig. 6A)**, with relative binding ratios indicating that a fraction of KIF1A associates with TDP-43 **(Fig. 6D)**. Reciprocal IP of KIF1A confirmed this interaction, co-purifying TDP-43 **(Fig. 6E, Supplementary Fig. 6B)**. The observed binding ratios likely reflect the relative abundance of these proteins and the stoichiometry of motor-cargo complexes.

**Figure 6.**
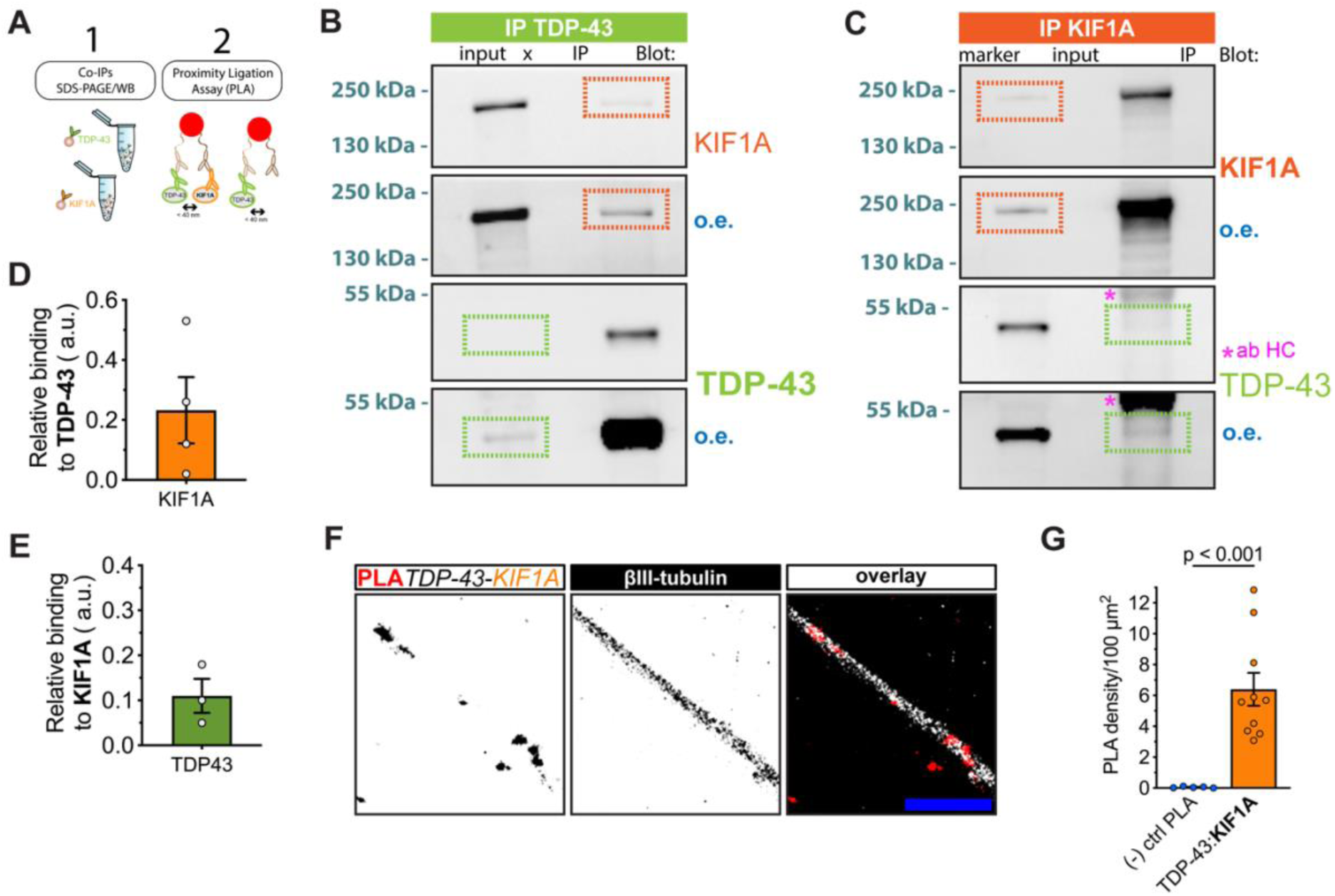
TDP-43 alternatively associates with KIF1A synaptic vesicle motor protein. **(A)** Schematic representation of co-IP and PLA experiments conducted at DIV 40 in human neurons using TDP-43 and KIF1A antibodies to test their interaction. **(B)** Representative immunoblots of TDP-43 IP bands (green box) and KIF1A co-IP bands (orange box). Each protein is represented by two images to show both unsaturated and over-exposed images (o.e.). **(C)** Representative immunoblots of KIF1A IP bands (orange box) and TDP-43 co-IP bands (green box). Each protein is represented by two images to show both unsaturated and over-exposed images (o.e.). **(D)** Co-IPs ratios expressed as KIF1A intensity relative to TDP-43 IP (*n*=4 biological replicates). **(E)** Co-IPs ratios expressed as TDP-43intensity relative to KIF1A IP (*n*=3 biological replicates). **(F)** Representative micrographs of the PLA experiments performed to validate the interaction of TDP-43 with KIF1A. Images show (+) PLA signal (red), and the axonal marker βIII-tubulin (white), and the overlay of the two channels highlighting the red spots distribution (scale bar = 10 µm). **(G)** Quantification of PLA densities in TDP-43:KIF1A *vs* TDP-43-(-)ctrl (*n*=5 images and *n*=10 images from 3 different biological replicates for TDP-43-(-)ctrl PLAs and TDP-43:KIF1A, respectively). Data show mean ± S.D **(D, E, and G)**. Statistical comparison was performed using unpaired *t*-test with Welch’s correction. Approximate *p* values are shown in the figure.

On the other hand, PLA confirmed close spatial association between TDP-43 and KIF1A in neuronal projections. Using βIII-tubulin_GFP cytoskeletal tracker to mark neurites, confocal imaging revealed positive PLA puncta densities along neuronal projections **(Fig. 6F, Supplementary Fig. 6C)**. Quantification showed significantly higher PLA puncta density for TDP-43-KIF1A compared to negative controls **(Fig. 6G)**. This validates the biochemical interaction data and demonstrates that TDP-43 and KIF1A associate within the axonal compartment.

Collectively, these findings reveal that TDP-43 engages with multiple anterograde motor systems, both kinesin-1 (KIF5A/B/C via KLC1) and kinesin-3 (KIF1A). This dual motor engagement may provide functional flexibility for TDP-43-containing MLOs transport regulation.

## Discussion

TDP-43 was initially defined as a predominantly nuclear protein, with its cytosolic presence considered primarily pathological. However, accumulating evidence indicates that TDP-43 physiologically localizes to the cytoplasm at lower concentrations than in the nucleus, including axonal and dendritic subdomains where it regulates mRNA transport and local translation^78,79^ TDP-43 axonal transport has been demonstrated in vivo in spinal motor neurons and other neuronal types using live imaging in Drosophila, zebrafish, and mouse models. Understanding the physiological mechanisms regulating TDP-43 axonal trafficking is therefore essential for elucidating how these processes become disrupted in ALS, where TDP-43 pathology is present in ∼97% of cases^80,81^.

It remains unclear how pathogenic TDP-43 spreads through neural connections to drive neurodegeneration in human ALS. Recent evidence reveals that axonal TDP-43 pathology is significantly increased in both upper motor neuron (UMN) axons within the descending corticospinal tracts (CST) and lower motor neuron (LMN) axons within the ventral horn gray matter of ALS patients compared to controls (Liu, Ong, Meng, et al., 2025 – submitted). These findings suggest shared molecular mechanisms underlying axonal TDP-43 pathology across the three clinical forms of ALS UMN-predominant, LMN-predominant, and typical ALS. The presence of TDP-43 pathology in both UMN and LMN axons supports the concept of a common pathogenic process affecting both motor neuron populations.

Recently, Tsuboguchi et al. demonstrated that mutant TDP-43 can spread along neuro-glial connections in the corticospinal pathway and induce distinct types of degeneration in spinal circuits *in vivo*^82^. Using adeno-associated virus (AAV) to express mutant TDP-43 in mice, they found that TDP-43 expressed in UMNs moved anterogradely along corticospinal tract axons and was transferred to oligodendrocytes, resulting in mild axonal degeneration. In contrast, TDP-43 expressed in LMNs did not spread retrogradely but caused severe motor neuron loss and degeneration of neighboring spinal neurons, leading to muscle atrophy. Collectively, studies in both human ALS and mouse models demonstrate that pathological TDP-43 can spread anterogradely from cortical neurons along their axons, coinciding with axonal degeneration and contributing to ALS progression.

Consistent with these findings, our observation that axonal TDP-43 transport can occur via alternative motor proteins shows a directional bias that may favor a cortical origin in human H9-derived cortical neurons. Together with ALS autopsy findings (Liu, Ong, Meng, et al., 2025, submitted), these results further support the notion that TDP-43 can be transported within the corticospinal tract (UMN axons) and may influence LMNs.

To understand the physiological mechanisms that may become disrupted in disease, we characterized TDP-43 axonal transport by comparing its dynamics to those of well-established cargoes with defined motor dependencies: Rab5 (predominantly retrograde, dynein-mediated)^56^, APP (bidirectional with slight anterograde bias, kinesin-1/dynein)^59^, and SYP (strongly anterograde, KIF1A-mediated)^58^. This comparative framework allowed us to infer which motor protein families likely regulate TDP-43 trafficking, particularly during its anterograde journey toward presynaptic terminals, since dynein–dynactin is the primary motor mediating retrograde transport for most cargoes^63^.

Our net transport analysis revealed that TDP-43 MLOs exhibit balanced bidirectional movement, with nearly equal proportions of anterograde and retrograde transport. This feature, also observed for other mRNA-localizing RBPs^54,83^, distinguishes TDP-43 from vesicular-bound cargoes that display stronger directional bias. Such balanced dynamics suggest that TDP-43 granules deliver mRNA cargoes to multiple axonal regions rather than a single subdomain, consistent with TDP-43’s role in maintaining diverse aspects of RNA axonal homeostasis^14,84,85^.

Despite this bidirectional character, TDP-43 displayed fast and processive dynamics similar to APP and SYP, rather than the slower, pause-prone Rab5. TDP-43 exhibited higher reversals, but fewer pauses, and traveled distances comparable to vesicular-bound cargoes. Notably, TDP-43 total track lengths were comparable to SYP and Rab5, but significantly lower than APP, which showed the widest range of traveled distances. This suggests TDP-43 transport encompasses both short- and long-range movements, consistent with the requirement for local mRNA delivery and translation at diverse axonal locations.

Velocity distribution analysis provided further mechanistic insight. TDP-43 anterograde velocities showed near-complete overlap with APP, strongly suggesting shared motor protein involvement. The positive correlation between segment duration and velocity for TDP-43 anterograde movement indicates processive motor engagement, a characteristic feature of kinesin-mediated transport. In contrast, TDP-43 retrograde velocities were distinct from all reference vesicles, suggesting that while dynein-dynactin generally mediates retrograde movement for most cargoes, it exhibits protein-specific processivity regulation^64,66,70^, likely through differential engagement with adaptor proteins. This unique retrograde profile may reflect specialized requirements for returning RNA granules to the soma.

Co-immunoprecipitation and proximity ligation assays revealed that TDP-43 associates with both KLC1 and DCTN1, consistent with its bidirectional transport profile. KLC1 knockdown markedly reduced TDP-43–KLC1 association, confirming interaction specificity. Remarkably, TDP-43 showed positive interaction with all three-mammalian kinesin-1 isoforms, KIF5A, KIF5B, and KIF5C^86,87^. This broad engagement may provide functional flexibility, allowing TDP-43 to use distinct transport pathways depending on neuronal compartment or cargo composition. This model gained further support when we observed that TDP-43 also associates with KIF1A, a kinesin-3 family member best known for transporting synaptic vesicle precursors^58,88^. This finding was consistent with our observation that TDP-43 shares transport characteristics with SYP, including processive anterograde runs. The engagement of TDP-43 with both kinesin-1 and kinesin-3 families suggests a multi-motor transport mechanism that may serve distinct functional roles. KIF1A could facilitate rapid, long-distance delivery of specific TDP-43 granule populations to distal synaptic terminals, while kinesin-1 may mediate more regulated transport or delivery to intermediate axonal regions. Alternatively, distinct TDP-43-containing MLOs, defined by their mRNA cargoes or phase-separation properties, may selectively recruit different motor systems.

An important implication of our findings is that TDP-43 MLOs display transport dynamics comparable to classical vesicular cargoes. Typical models of axonal transport have been predominantly developed from vesicular cargo studies, where transmembrane proteins serve as recognition sites for motor-adaptor complexes^29,32^. The fact that TDP-43 MLOs use similar motor systems raises key mechanistic questions: How do motors recognize and bind to these condensates in the absence of membranes? Are specific RNA sequences or RBP motifs serving as adaptor recruitment sites? Does the biophysical state of TDP-43 condensates modulate motor engagement? While previous studies proposed that RNA granules ’hitchhike’ on motile organelles such as lysosomes or mitochondria^32,33,89^, our results demonstrate that TDP-43 can also closely associate with motor proteins. This suggests that TDP-43 complexes possess an intrinsic motor-binding capacity independent of membrane-bound organelles, similar to other RNA binding proteins^54^, and regulatory microtubule-binding proteins known to engage motor proteins for mRNA transport^50^. TDP-43 axonal transport has been also demonstrated *in vivo*^21,90^ in spinal motor neurons and other neuronal types. However, whether both direct motor engagement and hitchhiking mechanisms operate for TDP-43-bound mRNA delivery to distinct axonal subdomains remains to be investigated. Moreover, whether TDP-43 transport is differentially regulated between UMNs and LMNs, both of which undergo axonal degeneration in ALS (Liu, Ong, Meng, et al. 2025 – submitted) but may exhibit distinct vulnerabilities, it is currently unknown and needs future investigation.

Although these experiments were not performed in neurons from ALS patients, these findings provide insight into mechanisms affected in ALS. Disrupted anterograde delivery of TDP-43 to distal axons compromises local mRNA translation essential for synaptic function and axonal maintenance^18,19,22,23^. Studies in familial ALS models with *Tardbp* mutations or *C9orf72* repeat expansions demonstrate defects in mRNA trafficking and local translation, supporting this notion^6,67,91^. Moreover, impaired retrograde transport may hinder clearance of TDP-43 granules from axons, promoting cytoplasmic aggregation. Consistent with this, mutations in dynein–dynactin components have also been identified and linked to ALS and other motor neuron diseases^66,92,93^. Pathological TDP-43 aggregates could sequester motor proteins or adaptors, further impairing axonal transport^13,94,95^. The involvement of multiple motor systems could provide redundancy, allowing partial compensation if one pathway fails (e.g., KIF5A dysfunction which is known to lead to ALS^96,97^). However, combined impairments from oxidative stress, mitochondrial failure, or cytoskeletal disruption, all common in ALS, could collapse this finely tuned and coordinated activities. Understanding these physiological mechanisms in future studies, perhaps using motor neurons derived from iPSC lines from ALS patients^98,99^, could provide a foundation for dissecting how TDP-43 transport becomes perturbed in ALS and may reveal therapeutic targets for restoring normal TDP-43 localization and function.

These findings reveal that TDP-43 utilizes a multi-motor transport system under physiological conditions. In disease, simultaneous disruption of multiple transport pathways could contribute to progressive accumulation of non-functional TDP-43 granules, impairing mRNA trafficking and local translation, and ultimately exacerbating neuronal vulnerability. Our results provide a mechanistic foundation for understanding how different physiological TDP-43 transport pathways can be dysregulated in ALS and highlight axonal TDP-43 and its trafficking pathways as potential therapeutic targets. Combined with evidence of glial activation, inflammation, myelin loss in the CST, and LMN degeneration in the ventral horn, our results together with those showed by Liu, Ong, Meng, et al. suggest that pathogenic TDP-43–related axonal transport dysfunction can drive motor system degeneration in ALS through multiple, interconnected mechanisms.

## Methods

### Differentiation of human neurons from an embryonic stem cell line

Human Neural Stem Cells (hNSCs) derived from the NIH approved H9 (WA09) human embryonic stem cell line was purchased from Merck (Germany). The hNSCs were plated on 100 mm petri dishes pre-coated with Matrigel (Corning) for 1 h. Cells where then maintained in culture with NSCs expansion media (KO DMEM/F12, 2% StemPro Neural Supplement, 1% Glutamax, 20 ng/ml β-FGF, 20 ng/ml EGF), which was exchanged every second day (DIV0-3). Upon reaching confluency, we. Performed a complete media change to favor neural progenitors’ induction (DMEM/F12, 1% B27, 0.5% N2, 1% Glutamax) by feeding the cells every other day until DIV 9. Neural progenitors were then detached with accutase (Thermo Fisher), centrifuged at 300xg for 5 min at RT, and the pellet was resuspended in Neuronal Optimized Media complete (NOMc) (DMEM-F12, 2% B27, 1% N2, 1 µg/ml laminin, 100 nM cAMP, 200 ng/ml ascorbic acid, 10 ng/ml BDNF, 10 ng/ml GDNF, 10 ng/ml IGF). Cells were counted and then seeded at different densities depending on the experimental needs. Neurons were cultured up to DIV 40 by feeding them with NOMc every 6 days.

### Plasmid design and preparation for the over-expression of fluorescently tagged proteins for axonal transport studies

Plasmids over-expressing TDP-43, Rab5, SYP, and APP were designed in a pcDNA 3.1(+) backbone, under a CMV promoter and GFP tag placed for all the constructs in the C-terminal region, except for the Rab5 where the fluorescent tag was placed at the N-terminal. The design was then transferred to a company (GenScript Biotech, New Jersey, US), which manufactured the stock plasmids. DHα bacterial competent cells (Thermo Fisher) were transformed by performing the following protocol: i) bacterial cells were retrieved from – 80° C and slowly thawed on ice, then the desired amount of stock DNA relative to each plasmid was incubated with the DHα competent cells; ii) cells and DNA were incubated for 30 min on ice, then the mix exposed for 1 min at 42° C, and finally again on ice for 5 min; iii) after that S.O.C. medium (Super optimal broth with Catabolite repression – Merck-Sigma – made of 2% tryptone, 0.5% yeast extract, 10 mM NaCl, 2.5 mM KCl, 10 mM MgCl_2_, 10 mM MgSO_4_, and 20 mM glucose) was added to the mix to reach 500 µl of total volume, and the mix was incubated for 1 h at 37° C shaking at 250 rpm; iv) after the incubation cells were centrifuged a 5000xg for 3 min and almost all the supernatant was discarded except for 50 µl used for pellet resuspension; v) mixed samples were plated independently on 100 mm petri dishes completed with LB agar containing the ampicillin to guarantee the selective growth of transformed bacteria. After 24 h of incubation up to 3 colonies per plasmid were selected and DNA was purified to be screened by gel electrophoresis. Gels were prepared with 2% agar to test for the correct size of the inserted plasmid. Next, in-gel DNA digestion was performed and the purified DNA was sent for sequencing analysis of the ORFs of interest. At last, the colonies selected per each plasmid were amplified to generate a long term transformed bacteria stock stored at -80°C. The cells were grown into a liquid LB medium containing ampicillin (1µg/µl). On the second day, we performed a maxi prep DNA isolation according to the Qiafilter Plasmid Maxi kit (Qiagen) protocol with which DNA was purified. Plasmids were aliquoted for long term at -20° C, and transfections were performed always with the same mother DNA samples.

### Plasmids transfection approach

All plasmids were transfected using the same protocol and strategy. 48 h before live imaging of fluorescently tagged protein, neurons were transfected with a 1:1 ratio of Lipofectamine 2000 (Thermo Fisher) and 1µg of DNA of interest:

- pcDNA3.1(+)-TDP-43_GFP
- pcDNA3.1(+)-GFP_Rab5
- pcDNA3.1(+)-Synaptophysin_GFP
- pcDNA3.1(+)-APP_GFP

For the ibidi multichannel slide, we used 50 µl of transfection mix per channel, for a total of 300 µl of the mix for all 6 channels. The transfection mix was prepared according to the manufacturer’s suggestion. In brief, 2 different tubes were prepared to incubate separately at R.T. for 5 min both Lipofectamine 2000 and the DNA of interest using NOMc. Then the two mixes were combined to form a final transfection mix, which was incubated for 20 min at R.T. prior to being added to neurons. The solution was kept in cell cultures for 2 h at 37° C and 5% CO_2_. After transfection, the Lipofectamine-containing solution was completely replaced with fresh NOMc, and cells were examined under a fluorescent microscope 24 h post-transfection.

### Lentiviral vectors transduction for protein-protein interaction experiments

Lentiviral constructs LV_shRNA-KLC1_GFP and LV_shRNA-scramble_GFP were designed for expression under the human synapsin 1 (hSyn1) promoter and were cloned and packaged by Origene (Rockville, MD, US). Human neurons (hNeurons) were transduced with lentiviral particles at DIV 16. Particles were retrieved from -80°C and slowly thawed on ice for 20 min. Lentiviral particles were then resuspended in NOMc media according to the validated M.O.I. and TU/ml provided by the manufacturer. Transduction was performed by replacing the culture media with NOMc-containing lentiviral particles for approximately 24 h. The following day, the lentivirus-containing media was completely replaced with fresh one. Transduction efficiency was monitored every 48 h until DIV 40.

### Live imaging and tracking

Live-cell imaging recordings were performed at 4fps (total = 240 frames) using a confocal microscope equipped with a live module (Zeiss Confocal LSM780, Zeiss Live LSM7) and an immersion oil objective 63X/1.4 NA Plan Apochromat. All the cell cultures were incubated with fresh artificial cerebrospinal fluid (aCSF - NaCl 121 mM, KCl 2.5 mM, CaCl_2_ 2.2 mM, MgSO_4_ 1 mM, NaHCO_3_ 29 mM, NaH_2_PO_4_ 0.45 mM, Na_2_HPO_4_ 0.5 mM, Glucose 20 mM) at pH 7.4, before starting live imaging experiments.

Time-lapse movies were processed with ImageJ before the analysis in Imaris (version 9.2, Oxford Instruments). The latter calculates a variety of movement parameters starting from a reference point. This starting point is established during the pre-processing in Image J where all the axons were transformed to be oriented with the cell body on the left and axon terminal on the right, hence, to track the anterograde movement from left to right and retrograde movement from right to left (**Supplementary Movie 1,2,3, and 4**). This pre-processing step is crucial for establishing the proportions of vesicles moving in anterograde or retrograde direction. Vesicles were segmented and tracked with the semi-automated spot tracking tool by applying an Autoregressive Motion algorithm. The algorithm required the input of the following parameters related mainly to the size of the vesicles and information about the continuity of movement over-time: XY estimated diameter, max distance, and max gap size. The diameter was chosen on an average empirical value for each specific cargo analyzed. For either max distance or max gap size, both spatial and temporal resolutions of the acquired movies were considered (temporal resolution = 4 fps; spatial resolution with the 63X objective described earlier in the paragraph = 0.104µm/pixel). Among all the statistical values obtained, those used for analyses of axonal transport parameters were the spatial displacement (ΔDx (t1, t0) = Px(t1) – Px(t2)) of the particles in each frame and the track duration (td=total time during which a particle moves). For transport dynamics analysis, we first divided the tracks into stationary or moving. All the tracks moving at < 10 s were excluded. For the net axonal transport analysis, tracks with a total track displacement in x < 250 µm were considered completely stationary. All the tracks with ≥ 0.1 µm/s velocity were considered motile and classified as anterograde or retrograde based on the directionality of movement. In the segmental axonal transport analysis, we used the Δ*x* displacement between frames to compute instantaneous changes in movement, which gave us more mechanistic information about cargoes features, like real-time movement ratios, reversal and pause frequencies, segmental velocities/run lengths. Finally, since the software computes the distance traveled by each particle along the *x* and *y* vectors, these values were presented as track lengths.

### Immunocytochemistry (ICC)

Neurons differentiated either in microfluidic or ibidi chambers (6 channels or 8 multi well-8mw) were fixed in 4% paraformaldehyde (PFA)/ 4% Sucrose for 1 h or 40 min, respectively. Incubation with 0.1 M Glycine for 5 min was used to quench the fixation. After that, cells were permeabilized with 0.1% Triton X-100 for 10 min. Samples were blocked with 5% BSA for 30 min and then incubated overnight at 4°C with primary antibodies in 3% BSA. The second day, the cells were incubated with secondary antibodies in 3% BSA for 2 h. Finally, cells were stained with DAPI for 3 min, then washed with ddH_2_O and mounted either with 50% Glycerol in 0.01% Na azide/PBS (ibidi 6 channels) or with Mowiol (ibidi 8mw and coverslips). Samples were then dried and stored at 4°C.

### Identification of axonal projections in a non-microfluidic chamber system

Studies relative to TDP-43, Rab5, SYP, and APP movement were carried in ibidi µ-Slides VI 0.4. Differently from the microfluidic chambers, the ibidi device lacks a physical division between the somato-dendritic and axonal compartments. For this reason, before seeding the neurons, the ibidi were marked with an arbitrary reference point to set an X;Y (0;0), this was then matched with the 0;0 of the confocal stage before time-lapse acquisition^44^. While acquiring movies, each selected projection was marked into a position list and saved. Post-live imaging cells were used for ICC, following the protocol described above, and positions were retrieved to identify pNFH(+) neurites with overlapping GFP(+) intensities, which were then included in the axonal transport analysis.

### Image analysis

Images acquired following ICC were analyzed either with ImageJ or Imaris.

1. <u>pNFH/GFP post-live imaging</u>: neurites were traced in ImageJ following GFP intensities for all the experiments that involved live imaging. Intensity profiles of TDP-43, Rab5, SYP, and APP conjugated to the GFP tag were matched with those of pNFH to either include or exclude the neurite from further axonal transport analysis^44^.
2. <u>Proximity Ligation Assay for protein-protein interaction</u>: z-stacks acquired at 40X or 63X magnification were analyzed to quantify the amount of PLA (+) dots, which indicate close interaction between two proteins of interest, and the frequency of the interaction within a certain distance. Images were masked for axonal markers, either βIII-tubulin and pNFH alone, or pNFH in combination with GFP, in the case of neuronal cell cultures transduced with shRNA-KLC1_GFP or scramble_GFP control. PLA dots along the axonal projections were measured using the surface/spots package of Imaris. On the other hand, the frequency of interaction was analyzed by tracing manually 10 axonal projections in each z-stack and then, with the Analyze particles tool from ImageJ we counted the number of PLA(+) dots along the projections.

### Proximity Ligation Assay (PLA) for protein-protein interaction studies

To confirm protein-protein interactions with an alternative approach, besides the co-IPs experiments, we performed a series of PLAs, establishing a close interaction between two proteins that are less than 40 nm apart in their physiological environment.

In this study, we performed PLA to detect interactions between the following protein pairs:

- TDP-43: KLC1
- TDP-43: KIF5A
- TDP-43: KIF5B
- TDP-43: KIF5C
- TDP-43: KIF1A

In addition, we also performed PLAs for both technical and biological controls:

- Technical (-): 1 antibody with PLA buffers and enzymes
- Technical (+): 2 different antibodies recognizing the same protein in different epitopes

For the biological controls were selected pairs of proteins known to be interacting/non-interacting:

- KLC1:KHC (kinesin heavy chain – constitutively interacting)
- KLC1:MyoV (myosin V – not interacting)
- TDP-43:hnRNPC (interacting in the nucleus)
- TDP-43:UBQLN2 (interacting in the cytosol)

The first part of a PLA protocol follows the same steps of an ICC, up to the incubation with primary antibodies (for further details see the previous paragraph on ICC). For the second part of the protocol, we followed the manufacturer’s instructions and used buffers, enzymes, and probes provided in the kit (DuoLink PLA, Merck). Briefly, per each reaction performed in an 8 mw ibidi chamber, we used 10 µl of both (+) and (-) probes, each corresponding to the host species of the primary antibodies used in the first step (usually anti-rabbit PLUS, and either anti-mouse or anti-goat MINUS). The solution was prepared in Ab diluent in a final volume of 100 µl. After removing the primary antibodies mix, the probe solution was incubated with the sample for 1 h at 37°C in a humid chamber. In the meantime, a ligation buffer and ligase enzyme were mixed. After removing the probes, the ligase mix was incubated for 30 min at 37°C with the sample. During this step, the amplification mix was prepared by adding in the solution the amplification buffer and the polymerase (0.5 µl/PLA reaction) in a final 100 µl volume.

The ligase mix was then removed and exchanged with the polymerase solution, which was incubated for 100 min at 37°C. After this step, additional secondary antibodies conjugated with fluorophores can be incubated to complete the PLA and the ICC of protein markers not involved directly in the assay. After the canonical final washes in PBS and the DAPI staining, samples were mounted in a 50/50 Glycerol/pure water solution and stored at 4°C.

### Protein extraction

For Immunoprecipitations (IPs), proteins from 40 DIV differentiated hNeurons, were collected using IP lysis buffer (1% Nonidet – P 40, 25 mM Tris buffer pH 7.4, 150 mM NaCl, 1mM EDTA, 5% Glycerol). Cells were incubated for 30 min on ice and cell membranes were disrupted using an insulin syringe. Supernatants were collected into new tubes following centrifugation for 20 min at 20000xg 4°C. Total lysates were quantified using the BCA assay according to the manufacturer’s protocol (Pierce™ BCA Protein Assay Kit, Thermo Fisher).

### Immunoprecipitations of endogenous bait proteins

Bait proteins were immunoprecipitated with Dynabeads Protein G (Thermo Fisher), which allows magnetic separation of immunocomplexes.

For IP of TDP-43, KLC1, DCTN1, KIF1A, and IgG isotype controls 50 µl of Dynabeads Protein G were used per each sample. Beads were vortexed for 30 seconds and then placed into DynaMag-2 to discard the storage solution. Then, the antibody conjugation step was performed by adding 200 µl of the Ab Binding Buffer (Thermo Fisher), and 5 µg of primary antibody specific for each protein (**Supplementary Table 1**). The antibody mix was incubated for 1 h at 4°C rotating end-over-end. After discarding the Ab binding buffer 500 µg of total lysate was added to the beads and samples incubated for 2 h at 4°C rotating end-over-end. Tubes were spun at 1000xg for 1 min and then placed into DynaMag-2 (Thermo Fisher) for separation of flow-through fractions. The beads were washed twice for 2 min with 200µl of wash buffer (150 mM NaCl, 50 mM Tris/Cl pH 7.5). Protein complexes were eluted using 1X LDS sample buffer, 1X NuPage DTT reducing agent (Thermo Fisher) and heated at max 80°C for 15 min. Eluates were transferred into new tubes to be used for SDS-PAGE and western blot analyses. Quantification of bound fractions to the respective bait proteins was performed by analyzing the ratio of band intensities in the co-IP complexes. The formula used was:

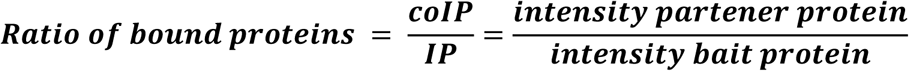

### SDS-PAGE and Western blotting

Quantified protein samples were loaded into Bolt Bis-Tris Plus 4-12 % precast gels (Novex, Thermo Fisher) and SDS-PAGE performed first for 10 min at 50 V and then at 100 V for 1 h 30 min. Proteins were then transferred onto PVDF 0.45 µm membranes (Thermo FIsher) using a Mini blot module (Invitrogen) for 1 h 50 min at 30 V. Membranes were washed 3X 5 min with 20mM Tris-Buffer (TBS) and blocked in 5% non-fat dry milk (NFDM) in 20mM TBS/0.2% Tween-20 (TBS-T) for 1 h at RT. After 3 washes in TBS, the membranes were probed overnight at 4° C with primary antibodies (**Supplementary Table 1**) in 1% BSA/TBS-T 0.2%. The second day, HRP-conjugated antibodies were prepared in 1% NFDM TBS-T 0.2% and incubated for 2 h at 4°C. Last, the membranes were washed 3X 5 min with TBS-T 0.2% and proteins detected using Chemiluminescent Substrate (SuperSignal™ West Pico PLUS, Thermo Scientific) by acquiring images in a Chemidoc (BioRad).

### Statistical analysis and data representation

All analyses were performed using Prism 10.0 (GraphPad Software). Data for axonal transport parameter determination were initially processed in Microsoft Excel tables, where multiple formulas were applied to calculate axonal transport parameters (e.g., real-time movement, reversal and pause frequency, segmental velocity). Statistical analyses and *post hoc* tests for multiple comparisons are described in detail in the figure legends, as well as the data representation for each graph shown in the corresponding figure. All experiments were performed at least three times using independent biological samples. The statistical pooling of data was performed depending on the biological message that the graph wanted to show for that specific parameter, particularly in the case of axonal transport analyses (net or segmental approach). For axonal transport parameters the results are represented as SuperPlots^45^ showing the individual data points in different colors, superimposed with the data points of either the biological replicate (net axonal transport analysis) or the grouped axons within that replicate. Regardless of grouping level, the mean of each experiment is shown as mean ± S.D.

The statistical analysis included unpaired *t*-test, unpaired *t*-test with Welch’s correction, One-way ANOVA, and the Kruskal-Wallis test. Proportional parameters such as anterograde *vs* retrograde were compared within each protein using unpaired *t*-test. In the case of PLA experiments when comparing KIF5A-, KIF5B-, KIF5C-, and KIF1A-TDP-43 interactions *vs* control, unpaired *t*-test with Welch’s correction was used. When comparing the means of three or more normally distributed samples such as for time in motion or track density and post KLC1 shRNA PLA we performed a one-way ANOVA followed by Dunnett’s and Tukey’s multiple comparison tests, respectively, as specified in the figure legends. For data that did not follow a normal distribution such as reversal or pause frequency, track length, and frequency of segmental velocities, the Kruskal Wallis test was performed followed by Dunn’s *post hoc* test.

Correlations between segmental velocity and segment duration were assessed using Spearman’s rank correlation coefficient (ρ). To evaluate whether the strength of the correlation differed between anterograde and retrograde transport, correlation coefficients were Fisher’s z-transformed and compared using a two-tailed z-test.

No statistical methods were used to predetermine sample size; however, sample sizes are comparable to those reported in previous axonal transport publications^46–48^. As described early in the paragraph multiple independent experiments were performed using multiple sample replicates as detailed in the figure legends.

## Supplementary Information

Supplementary material will be available online.

## Data availability

All relevant data associated with the current study are available in the manuscript or the supplementary material. Additional information about data and protocols is available from the authors upon request.

## Acknowledgments

We thank all present and past members of the Stokin Lab (Translation and Neuroscience Aging Program laboratory) for their invaluable support throughout the development and completion of this study.

## Author contribution

M.F. and G.B.S. conceived the study, developed the methodology, and wrote the original draft. M.F. and V.M.P.D. performed the investigation, software development, formal analysis, and data curation. M.F., V.M.P.D., N.D., M.Č., and K.K. conducted experimental investigations, including co-immunoprecipitation assays and plasmid design and preparation. C.L.-S. supervised the study and contributed to manuscript editing. G.F., R.A.S., and C.N.S. provided supervision and funding acquisition and contributed to manuscript editing. G.B.S. managed project administration and funding acquisition. All authors reviewed the manuscript.

## Declaration

### Ethical approval

Not applicable.

## References

1 Ayala, Y. M. et al. TDP-43 regulates its mRNA levels through a negative feedback loop. EMBO J 30, 277–288 (2011). 10.1038/emboj.2010.310

2 Ayala, Y. M. et al. Structural determinants of the cellular localization and shuttling of TDP-43. J Cell Sci 121, 3778–3785 (2008). 10.1242/jcs.038950

3 Buratti, E. & Baralle, F. E. Characterization and Functional Implications of the RNA Binding Properties of Nuclear Factor TDP-43, a Novel Splicing Regulator ofCFTR Exon 9. Journal of Biological Chemistry 276, 36337–36343 (2001). 10.1074/jbc.M104236200

4 Chen-Plotkin, A. S., Lee, V. M. Y. & Trojanowski, J. Q. TAR DNA-binding protein 43 in neurodegenerative disease. Nature Reviews Neurology 6, 211–220 (2010). 10.1038/nrneurol.2010.18

5 Tollervey, J. R. et al. Characterizing the RNA targets and position-dependent splicing regulation by TDP-43. Nat Neurosci 14, 452–458 (2011). 10.1038/nn.2778

6 Altman, T. et al. Axonal TDP-43 condensates drive neuromuscular junction disruption through inhibition of local synthesis of nuclear encoded mitochondrial proteins. Nat Commun 12, 6914 (2021). 10.1038/s41467-021-27221-8

7 Briese, M. et al. Loss of Tdp-43 disrupts the axonal transcriptome of motoneurons accompanied by impaired axonal translation and mitochondria function. Acta Neuropathol Commun 8, 116 (2020). 10.1186/s40478-020-00987-6

8 Piol, D., Robberechts, T. & Da Cruz, S. Lost in local translation: TDP-43 and FUS in axonal/neuromuscular junction maintenance and dysregulation in amyotrophic lateral sclerosis. Neuron 111, 1355–1380 (2023). 10.1016/j.neuron.2023.02.028

9 Agra Almeida Quadros, A. R., et al. Cryptic splicing of stathmin-2 and UNC13A mRNAs is a pathological hallmark of TDP-43-associated Alzheimer’s disease. Acta Neuropathol 147, 9 (2024). 10.1007/s00401-023-02655-0

10 Nguyen, T. B. et al. Aberrant splicing in Huntington’s disease accompanies disrupted TDP-43 activity and altered m6A RNA modification. Nat Neurosci 28, 280–292 (2025). 10.1038/s41593-024-01850-w

11 Afroz, T. et al. Functional and dynamic polymerization of the ALS-linked protein TDP-43 antagonizes its pathologic aggregation. Nat Commun 8, 45 (2017). 10.1038/s41467-017-00062-0

12 Ravits, J. et al. Deciphering amyotrophic lateral sclerosis: what phenotype, neuropathology and genetics are telling us about pathogenesis. Amyotroph Lateral Scler Frontotemporal Degener 14 **Suppl 1**, 5–18 (2013). 10.3109/21678421.2013.778548

13 Ionescu, A., Altman, T. & Perlson, E. Looking for answers far away from the soma-the (un)known axonal functions of TDP-43, and their contribution to early NMJ disruption in ALS. Mol Neurodegener 18, 35 (2023). 10.1186/s13024-023-00623-6

14 Melamed, Z. et al. Premature polyadenylation-mediated loss of stathmin-2 is a hallmark of TDP-43-dependent neurodegeneration. Nat Neurosci 22, 180–190 (2019). 10.1038/s41593-018-0293-z

15 Klim, J. R. et al. ALS-implicated protein TDP-43 sustains levels of STMN2, a mediator of motor neuron growth and repair. Nat Neurosci 22, 167–179 (2019). 10.1038/s41593-018-0300-4

16 Brown, A. L. et al. TDP-43 loss and ALS-risk SNPs drive mis-splicing and depletion of UNC13A. Nature 603, 131–137 (2022). 10.1038/s41586-022-04436-3

17 Oberstadt, M., Classen, J., Arendt, T. & Holzer, M. TDP-43 and Cytoskeletal Proteins in ALS. Mol Neurobiol 55, 3143–3151 (2018). 10.1007/s12035-017-0543-1

18 Jiang, T. et al. Amyotrophic lateral sclerosis mutant TDP-43 may cause synaptic dysfunction through altered dendritic spine function. Dis Model Mech 12 (2019). 10.1242/dmm.038109

19 Bak, A. N. et al. Cytoplasmic TDP-43 accumulation drives changes in C-bouton number and size in a mouse model of sporadic Amyotrophic Lateral Sclerosis. Mol Cell Neurosci 125, 103840 (2023). 10.1016/j.mcn.2023.103840

20 Wong, C. E. et al. TDP-43 proteinopathy impairs mRNP granule mediated postsynaptic translation and mRNA metabolism. Theranostics 11, 330–345 (2021). 10.7150/thno.51004

21 Alami, N. H. et al. Axonal transport of TDP-43 mRNA granules is impaired by ALS-causing mutations. Neuron 81, 536–543 (2014). 10.1016/j.neuron.2013.12.018

22 Ionescu, A. et al. Muscle-derived miR-126 regulates TDP-43 axonal local synthesis and NMJ integrity in ALS models. Nat Neurosci (2025). 10.1038/s41593-025-02062-6

23 Broadhead, M. J. et al. Synaptic expression of TAR-DNA-binding protein 43 in the mouse spinal cord determined using super-resolution microscopy. Front Mol Neurosci 16, 1027898 (2023). 10.3389/fnmol.2023.1027898

24 Nagano, S. et al. TDP-43 transports ribosomal protein mRNA to regulate axonal local translation in neuronal axons. Acta Neuropathol 140, 695–713 (2020). 10.1007/s00401-020-02205-y

25 Birsa, N., Bentham, M. P. & Fratta, P. Cytoplasmic functions of TDP-43 and FUS and their role in ALS. Semin Cell Dev Biol 99, 193–201 (2020). 10.1016/j.semcdb.2019.05.023

26 Buratti, E. Functional Significance of TDP-43 Mutations in Disease. Adv Genet 91, 1–53 (2015). 10.1016/bs.adgen.2015.07.001

27 Loughlin, F. E. & Wilce, J. A. TDP-43 and FUS-structural insights into RNA recognition and self-association. Curr Opin Struct Biol 59, 134–142 (2019). 10.1016/j.sbi.2019.07.012

28 Zacco, E., Martin, S. R., Thorogate, R. & Pastore, A. The RNA-Recognition Motifs of TAR DNA-Binding Protein 43 May Play a Role in the Aberrant Self-Assembly of the Protein. Front Mol Neurosci 11, 372 (2018). 10.3389/fnmol.2018.00372

29 Fernandopulle, M. S., Lippincott-Schwartz, J. & Ward, M. E. RNA transport and local translation in neurodevelopmental and neurodegenerative disease. Nat Neurosci 24, 622–632 (2021). 10.1038/s41593-020-00785-2

30 Sebastian Baumann, A. K., Maria Gili, Verena Ruprecht, Stefan Wieser, Sebastian P. Maurer. A reconstitued mammalian APC-kinesin complex selectively transports defined packages of axonal mRNAs. Science Advances (2020).

31 Wu, H., Zhou, J., Zhu, T., Cohen, I. & Dictenberg, J. A kinesin adapter directly mediates dendritic mRNA localization during neural development in mice. J Biol Chem 295, 6605–6628 (2020). 10.1074/jbc.RA118.005616

32 Liao, Y. C. et al. RNA Granules Hitchhike on Lysosomes for Long-Distance Transport, Using Annexin A11 as a Molecular Tether. Cell 179, 147–164 e120 (2019). 10.1016/j.cell.2019.08.050

33 Cioni, J. M. et al. Late Endosomes Act as mRNA Translation Platforms and Sustain Mitochondria in Axons. Cell 176, 56–72 e15 (2019). 10.1016/j.cell.2018.11.030

34 Buratti, E. TDP-43 post-translational modifications in health and disease. Expert Opin Ther Targets 22, 279–293 (2018). 10.1080/14728222.2018.1439923

35 Chen, Y. & Cohen, T. J. Aggregation of the nucleic acid-binding protein TDP-43 occurs via distinct routes that are coordinated with stress granule formation. J Biol Chem 294, 3696–3706 (2019). 10.1074/jbc.RA118.006351

36 Cohen, T. J. et al. An acetylation switch controls TDP-43 function and aggregation propensity. Nat Commun 6, 5845 (2015). 10.1038/ncomms6845

37 Brady, S. T. & Morfini, G. A. Regulation of motor proteins, axonal transport deficits and adult-onset neurodegenerative diseases. Neurobiol Dis 105, 273–282 (2017). 10.1016/j.nbd.2017.04.010

38 Dadon-Nachum, M., Melamed, E. & Offen, D. The “dying-back” phenomenon of motor neurons in ALS. J Mol Neurosci 43, 470–477 (2011). 10.1007/s12031-010-9467-1

39 Lepine, S., Castellanos-Montiel, M. J. & Durcan, T. M. TDP-43 dysregulation and neuromuscular junction disruption in amyotrophic lateral sclerosis. Transl Neurodegener 11, 56 (2022). 10.1186/s40035-022-00331-z

40 Hirokawa, N., Niwa, S. & Tanaka, Y. Molecular motors in neurons: transport mechanisms and roles in brain function, development, and disease. Neuron 68, 610–638 (2010). 10.1016/j.neuron.2010.09.039

41 McKenney, R. J., Huynh, W., Tanenbaum, M. E., Bhabha, G. & Vale, R. D. Activation of cytoplasmic dynein motility by dynactin-cargo adapter complexes. Science 345, 337–341 (2014). 10.1126/science.1254198

42 Elshenawy, M. M. et al. Cargo adaptors regulate stepping and force generation of mammalian dynein-dynactin. Nat Chem Biol 15, 1093–1101 (2019). 10.1038/s41589-019-0352-0

43 De Pace, R. et al. Messenger RNA transport on lysosomal vesicles maintains axonal mitochondrial homeostasis and prevents axonal degeneration. Nat Neurosci 27, 1087–1102 (2024). 10.1038/s41593-024-01619-1

44 Feole, M. et al. Swedish Alzheimer’s disease variant perturbs activity of retrograde molecular motors and causes widespread derangement of axonal transport pathways. J Biol Chem 300, 107137 (2024). 10.1016/j.jbc.2024.107137

45 Lord, S. J., Velle, K. B., Mullins, R. D. & Fritz-Laylin, L. K. SuperPlots: Communicating reproducibility and variability in cell biology. J Cell Biol 219 (2020). 10.1083/jcb.202001064

46 Maday, S., Twelvetrees, A. E., Moughamian, A. J. & Holzbaur, E. L. Axonal transport: cargo-specific mechanisms of motility and regulation. Neuron 84, 292–309 (2014). 10.1016/j.neuron.2014.10.019

47 Neumann, S., Chassefeyre, R., Campbell, G. E. & Encalada, S. E. KymoAnalyzer: a software tool for the quantitative analysis of intracellular transport in neurons. Traffic 18, 71–88 (2017). 10.1111/tra.12456

48 Reis, G. F. et al. Molecular motor function in axonal transport in vivo probed by genetic and computational analysis in Drosophila. Mol Biol Cell 23, 1700–1714 (2012). 10.1091/mbc.E11-11-0938

49 Lukavsky, P. J. et al. Molecular basis of UG-rich RNA recognition by the human splicing factor TDP-43. Nat Struct Mol Biol 20, 1443–1449 (2013). 10.1038/nsmb.2698

50 <A reconstitued mammalian APC-kinesin complex selectively transports defined packages of axonal mRNAs_Baumann_2020.pdf>.

51 Pozo Devoto, V. M., et al. Unraveling axonal mechanisms of traumatic brain injury. Acta Neuropathol Commun 10, 140 (2022). 10.1186/s40478-022-01414-8

52 Cason, S. E. & Holzbaur, E. L. F. Selective motor activation in organelle transport along axons. Nat Rev Mol Cell Biol 23, 699–714 (2022). 10.1038/s41580-022-00491-w

53 Ichinose, S., Ogawa, T. & Hirokawa, N. Mechanism of Activity-Dependent Cargo Loading via the Phosphorylation of KIF3A by PKA and CaMKIIa. Neuron 87, 1022–1035 (2015). 10.1016/j.neuron.2015.08.008

54 Fukuda, Y. et al. Binding and transport of SFPQ-RNA granules by KIF5A/KLC1 motors promotes axon survival. J Cell Biol 220 (2021). 10.1083/jcb.202005051

55 Deinhardt, K. et al. Rab5 and Rab7 control endocytic sorting along the axonal retrograde transport pathway. Neuron 52, 293–305 (2006). 10.1016/j.neuron.2006.08.018

56 Goto-Silva, L. et al. Retrograde transport of Akt by a neuronal Rab5-APPL1 endosome. Sci Rep 9, 2433 (2019). 10.1038/s41598-019-38637-0

57 Gordon, S. L. & Cousin, M. A. The Sybtraps: control of synaptobrevin traffic by synaptophysin, alpha-synuclein and AP-180. Traffic 15, 245–254 (2014). 10.1111/tra.12140

58 Kondo, M., Takei, Y. & Hirokawa, N. Motor protein KIF1A is essential for hippocampal synaptogenesis and learning enhancement in an enriched environment. Neuron 73, 743–757 (2012). 10.1016/j.neuron.2011.12.020

59 Chaves, R. S. et al. Amyloidogenic Processing of Amyloid Precursor Protein Drives Stretch-Induced Disruption of Axonal Transport in hiPSC-Derived Neurons. J Neurosci 41, 10034–10053 (2021). 10.1523/JNEUROSCI.2553-20.2021

60 Lazarov, O. et al. Impairments in fast axonal transport and motor neuron deficits in transgenic mice expressing familial Alzheimer’s disease-linked mutant presenilin 1. J Neurosci 27, 7011–7020 (2007). 10.1523/JNEUROSCI.4272-06.2007

61 Otero, M. G. et al. Proteasome stress leads to APP axonal transport defects by promoting its amyloidogenic processing in lysosomes. J Cell Sci 131 (2018). 10.1242/jcs.214536

62 Neumann, S., Campbell, G. E., Szpankowski, L., Goldstein, L. S. & Encalada, S. E. Characterizing the composition of molecular motors on moving axonal cargo using “cargo mapping” analysis. J Vis Exp, e52029 (2014). 10.3791/52029

63 Reck-Peterson, S. L., Redwine, W. B., Vale, R. D. & Carter, A. P. The cytoplasmic dynein transport machinery and its many cargoes. Nat Rev Mol Cell Biol 19, 382–398 (2018). 10.1038/s41580-018-0004-3

64 Canty, J. T. & Yildiz, A. Activation and Regulation of Cytoplasmic Dynein. Trends Biochem Sci 45, 440–453 (2020). 10.1016/j.tibs.2020.02.002

65 Cianfrocco, M. A., DeSantis, M. E., Leschziner, A. E. & Reck-Peterson, S. L. Mechanism and regulation of cytoplasmic dynein. Annu Rev Cell Dev Biol 31, 83–108 (2015). 10.1146/annurev-cellbio-100814-125438

66 Deshimaru, M. et al. DCTN1 Binds to TDP-43 and Regulates TDP-43 Aggregation. Int J Mol Sci 22 (2021). 10.3390/ijms22083985

67 Laszlo, Z. I. et al. Synaptic proteomics reveal distinct molecular signatures of cognitive change and C9ORF72 repeat expansion in the human ALS cortex. Acta Neuropathol Commun 10, 156 (2022). 10.1186/s40478-022-01455-z

68 Fu, M. M. & Holzbaur, E. L. JIP1 regulates the directionality of APP axonal transport by coordinating kinesin and dynein motors. J Cell Biol 202, 495–508 (2013). 10.1083/jcb.201302078

69 Fu, M. M. & Holzbaur, E. L. Integrated regulation of motor-driven organelle transport by scaffolding proteins. Trends Cell Biol 24, 564–574 (2014). 10.1016/j.tcb.2014.05.002

70 Ayloo, S. et al. Dynactin functions as both a dynamic tether and brake during dynein-driven motility. Nat Commun 5, 4807 (2014). 10.1038/ncomms5807

71 Chaaban, S. & Carter, A. P. Structure of dynein-dynactin on microtubules shows tandem adaptor binding. Nature 610, 212–216 (2022). 10.1038/s41586-022-05186-y

72 Tan, S. C., Scherer, J. & Vallee, R. B. Recruitment of dynein to late endosomes and lysosomes through light intermediate chains. Mol Biol Cell 22, 467–477 (2011). 10.1091/mbc.E10-02-0129

73 Cassel, J. A. & Reitz, A. B. Ubiquilin-2 (UBQLN2) binds with high affinity to the C-terminal region of TDP-43 and modulates TDP-43 levels in H4 cells: characterization of inhibition by nucleic acids and 4-aminoquinolines. Biochim Biophys Acta 1834, 964–971 (2013). 10.1016/j.bbapap.2013.03.020

74 Appocher, C. et al. Major hnRNP proteins act as general TDP-43 functional modifiers both in Drosophila and human neuronal cells. Nucleic Acids Res 45, 8026–8045 (2017). 10.1093/nar/gkx477

75 Hirokawa, N. & Tanaka, Y. Kinesin superfamily proteins (KIFs): Various functions and their relevance for important phenomena in life and diseases. Exp Cell Res 334, 16–25 (2015). 10.1016/j.yexcr.2015.02.016

76 Liao, P. et al. Association of variants in the KIF1A gene with amyotrophic lateral sclerosis. Transl Neurodegener 11, 46 (2022). 10.1186/s40035-022-00320-2

77 Hummel, J. J. A. & Hoogenraad, C. C. Specific KIF1A-adaptor interactions control selective cargo recognition. J Cell Biol 220 (2021). 10.1083/jcb.202105011

78 Fallini, C., Bassell, G. J. & Rossoll, W. The ALS disease protein TDP-43 is actively transported in motor neuron axons and regulates axon outgrowth. Hum Mol Genet 21, 3703–3718 (2012). 10.1093/hmg/dds205

79 Sephton, C. F. et al. Identification of neuronal RNA targets of TDP-43-containing ribonucleoprotein complexes. J Biol Chem 286, 1204–1215 (2011). 10.1074/jbc.M110.190884

80 Baxi, E. G. et al. Answer ALS, a large-scale resource for sporadic and familial ALS combining clinical and multi-omics data from induced pluripotent cell lines. Nat Neurosci 25, 226–237 (2022). 10.1038/s41593-021-01006-0

81 Prasad, A., Bharathi, V., Sivalingam, V., Girdhar, A. & Patel, B. K. Molecular Mechanisms of TDP-43 Misfolding and Pathology in Amyotrophic Lateral Sclerosis. Front Mol Neurosci 12, 25 (2019). 10.3389/fnmol.2019.00025

82 Tsuboguchi, S. et al. TDP-43 differentially propagates to induce antero- and retrograde degeneration in the corticospinal circuits in mouse focal ALS models. Acta Neuropathologica 146, 611–629 (2023). 10.1007/s00401-023-02615-8

83 Lopez-Erauskin, J. et al. ALS/FTD-Linked Mutation in FUS Suppresses Intra-axonal Protein Synthesis and Drives Disease Without Nuclear Loss-of-Function of FUS. Neuron 100, 816–830 e817 (2018). 10.1016/j.neuron.2018.09.044

84 Beccari, M. S. et al. Stathmin-2 enhances motor axon regeneration after injury independent of its binding to tubulin. Proc Natl Acad Sci U S A 122, e2502294122 (2025). 10.1073/pnas.2502294122

85 Lopez-Erauskin, J. et al. Stathmin-2 loss leads to neurofilament-dependent axonal collapse driving motor and sensory denervation. Nat Neurosci 27, 34–47 (2024). 10.1038/s41593-023-01496-0

86 Iworima, D. G., Pasqualotto, B. A. & Rintoul, G. L. Kif5 regulates mitochondrial movement, morphology, function and neuronal survival. Mol Cell Neurosci 72, 22–33 (2016). 10.1016/j.mcn.2015.12.014

87 Takemura, R. et al. mRNA expression of KIF1A, KIF1B, KIF2, KIF3A, KIF3B, KIF4, KIF5, and cytoplasmic dynein during axonal regeneration. J Neurosci 16, 31–35 (1996). 10.1523/JNEUROSCI.16-01-00031.1996

88 Hirokawa, N., Nitta, R. & Okada, Y. The mechanisms of kinesin motor motility: lessons from the monomeric motor KIF1A. Nat Rev Mol Cell Biol 10, 877–884 (2009). 10.1038/nrm2807

89 Shigeoka, T. et al. On-Site Ribosome Remodeling by Locally Synthesized Ribosomal Proteins in Axons. Cell Rep 29, 3605–3619 e3610 (2019). 10.1016/j.celrep.2019.11.025

90 Svahn, A. J. et al. Nucleo-cytoplasmic transport of TDP-43 studied in real time: impaired microglia function leads to axonal spreading of TDP-43 in degenerating motor neurons. Acta Neuropathol 136, 445–459 (2018). 10.1007/s00401-018-1875-2

91 Cook, C. N. et al. C9orf72 poly(GR) aggregation induces TDP-43 proteinopathy. Sci Transl Med 12 (2020). 10.1126/scitranslmed.abb3774

92 Sleigh, J. N. et al. Mice Carrying ALS Mutant TDP-43, but Not Mutant FUS, Display In Vivo Defects in Axonal Transport of Signaling Endosomes. Cell Rep 30, 3655–3662 e3652 (2020). 10.1016/j.celrep.2020.02.078

93 Sleigh, J. N., Rossor, A. M., Fellows, A. D., Tosolini, A. P. & Schiavo, G. Axonal transport and neurological disease. Nat Rev Neurol 15, 691–703 (2019). 10.1038/s41582-019-0257-2

94 Scialo, C. et al. Seeded aggregation of TDP-43 induces its loss of function and reveals early pathological signatures. Neuron 113, 1614–1628 e1611 (2025). 10.1016/j.neuron.2025.03.008

95 Vishal, S. S., Wijegunawardana, D., Salaikumaran, M. R. & Gopal, P. P. Sequence Determinants of TDP-43 Ribonucleoprotein Condensate Formation and Axonal Transport in Neurons. Front Cell Dev Biol 10, 876893 (2022). 10.3389/fcell.2022.876893

96 Nicolas, A. et al. Genome-wide Analyses Identify KIF5A as a Novel ALS Gene. Neuron 97, 1267–1288 (2018). 10.1016/j.neuron.2018.02.027

97 Baron, D. M. et al. ALS-associated KIF5A mutations abolish autoinhibition resulting in a toxic gain of function. Cell Rep 39, 110598 (2022). 10.1016/j.celrep.2022.110598

98 Workman, M. J. et al. Large-scale differentiation of iPSC-derived motor neurons from ALS and control subjects. Neuron 111, 1191–1204 e1195 (2023). 10.1016/j.neuron.2023.01.010

99 Ye, L. et al. Sporadic ALS hiPSC-derived motor neurons show axonal defects linked to altered axon guidance pathways. Neurobiol Dis 206, 106815 (2025). 10.1016/j.nbd.2025.106815

