## Supplementary data and figures for "Multiple motor proteins regulate ALS-linked TDP-43 anterograde axonal transport": Supplementary Information_Feole_et_al_2025.pdf

**Running title**: Regulation of TDP-43 anterograde axonal transport

**Keywords:** TDP-43; axonal transport; Amyotrophic Lateral Sclerosis; motor proteins; adaptors; cargoes.

### Supplementary Tables and Figures

#### Supplementary Table 1. Antibodies and constructs used in the study.

| Antibodies | Source | cat. No. |
|---|---|---|
| Mouse monoclonal anti-pNFH (SMI31) | BioLegend | 801601 |
| Rabbit monoclonal anti-p150 (Dynactin subunit 1) (D1W1O) | Cell Signaling | #69399 |
| Mouse monoclonal anti-p150 (Dynactin subunit 1) (12) | Santa Cruz Biotech. | sc-135890 |
| Mouse monoclonal anti-Dynein Intermediate Chain1 (DIC1) (74.1) | Merck | MAB1618 |
| Rabbit monoclonal anti-KLC1) (EPR12441(B)) | Abcam | 174273 |
| Mouse monoclonal anti-KLC1) (L2) | Santa Cruz Biotech. | sc-58776 |
| Rabbit polyclonal anti-KIF5A | Abcam | ab5628 |
| Rabbit monoclonal anti-KIF5B (EPR10276(B)) | Abcam | ab167429 |
| Rabbit monoclonal anti-KIF5C (EPR16224-32) | Abcam | ab193352 |
| Rabbit monoclonal anti-KIF1A (EPR11790(B)) | Abcam | ab180153 |
| Goat polyclonal anti- TDP-43 | Thermo Fisher | PA519064 |
| Mouse monoclonal anti-TDP-43 (GT733) | Thermo Fisher | MA527828 |
| Rabbit monoclonal anti-TDP-43 (EPR5810) | Abcam | ab109535 |
| Mouse monoclonal anti-Myosin Va (MyoVa) (G-4) | Santa Cruz Biotech. | sc-365986 |
| Mouse monoclonal anti-hnRNPC (4F4) | Santa Cruz Biotech. | sc-32308 |
| Rabbit polyclonal anti Ubiquilin2 (UBQLN2) | Novus Biologicals | 85639 |
| Mouse monoclonal anti-KHC (H2) | Merck | MAB1614 |
| Normal Mouse IgG | Santa Cruz Biotech. | sc-2025 |
| Normal Rabbit IgG | Novus biologicals | 81056910 |
| Rabbit polyclonal anti-β-III tubulin | BioLegend | 802001 |
| Mouse monoclonal anti-β-actin (AC-74) | Millipore-Sigma | A2228 |
| Alexa Fluor 488 donkey anti-mouse IgG | Thermo Fisher | A31570 |
| Alexa Fluor 488 donkey anti-rabbit IgG | Thermo Fisher | A21206 |
| Alexa Fluor 488 donkey anti-goat IgG | Thermo Fisher | A11055 |
| Alexa Fluor 546 donkey anti-mouse IgG | Thermo Fisher | A10036 |
| Alexa Fluor 546 donkey anti-rabbit IgG | Thermo Fisher | A10040 |
| Alexa Fluor 546 donkey anti-goat IgG | Thermo Fisher | A11056 |
| Alexa Fluor 555 donkey anti-mouse IgG | Thermo Fisher | A31570 |
| Alexa Fluor 555 donkey anti-rabbit IgG | Thermo Fisher | A31572 |
| Alexa Fluor 555 donkey anti-goat IgG | Thermo Fisher | A21432 |
| Alexa Fluor 647 donkey anti-mouse IgG | Thermo Fisher | A31571 |
| DyLight 647 donkey anti-mouse IgG | Abcam | ab150107 |
| Alexa Fluor 488 goat anti-guinea pig IgG | Abcam | ab150185 |
| Chicken anti-mouse IgG-HRP | Santa Cruz Biotech. | sc-2962 |
| Goat anti-Rabbit IgG-HRP | Cell Signaling | #7074 |
| Horse anti-Mouse IgG-HRP | Cell Signaling | #7076 |
| Mouse anti-Goat IgG-HRP | Santa Cruz Biotech. | sc-2354 |
| Clean-Blot$^{TM}$ IP detection reagent-HRP | Thermo Fisher | 21230 |
| **Constructs for over-expression of fluorescently tagged ORFs** | | |
| Bacmam 2.0 Tubulin_GFP | Thermo Fisher | C10509 |
| pcDNA3.1(+)-TDP-43_GFP | GenScript | ID: U4213DG120 |
| pcDNA3.1(+)-GFP_Rab5 | GenScript | ID: U282AEF060 |
| pcDNA3.1(+)-Synaptophysin_GFP | GenScript | ID: U282AEF060 |
| pcDNA3.1(+)-APP_GFP | GenScript | ID: U0869DK28 |

### Supplementary Figure 1

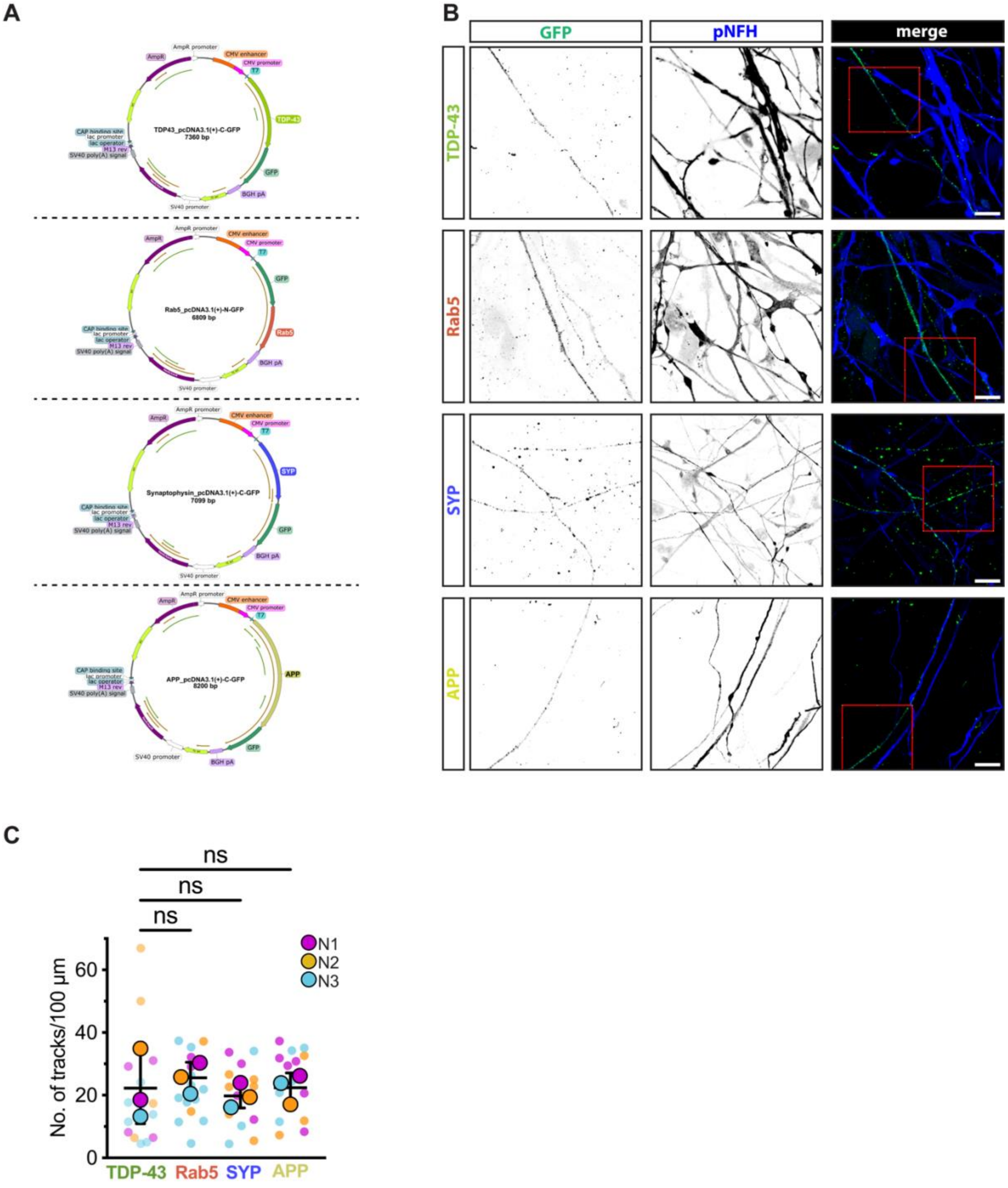

**Supplementary Figure 1 GFP_tag axonal cargoes TDP-43, Rab5, SYP, and APP are tracked in pNFH(+) neurites. (A)** Schematic maps of plasmid constructs used in this study. Plasmids were generated into pcDNA3.1(+) _GFP, and were cloned by inserting the human sequences of either TDP-43, Rab5, Synaptophysin, or APP under CMV promoter (Genscript). Key features are indicated in the SnapGene-generated maps. **(B)** Representative uncropped micrographs of neurons transfected with either TDP-43, Rab5, SYP, or APP_GFP (green), colocalizing with pNFH(+) (blue) axons. Red boxes indicate the crop areas reported in **Fig.1B** (scale bar = 20 µm). **(C)** Track density/100 µm representing the number of active trajectories detected along pNFH(+) neurites used for axonal transport analysis in Imaris. (*n* = 15 axons from *n* = 3 biological replicates). Data are represented as mean ± S.D. One-way ANOVA results are represented with Dunnet's pos-hoc multiple comparisons *vs* TDP-43 as the cargo of interest.

**Supplementary Table 2. Number of axons and tracks analyzed using Imaris software related to segmental analysis in Figure 2A, and B. Mean values of real time anterograde, retrograde and pausing, of tracks grouped per axon.**

| Axonal cargo | Nº of tracks (total) | Nº moving tracks | Real Time Anterograde (mean±S.D.) | Real Time Retrograde (mean±S.D.) | Real Time Pause (mean±S.D.) |
|---|---|---|---|---|---|
| TDP-43 | 366 | 266 | 37.23±7.72 | 36.42±7.41 | 26.34±14.02 |
| Rab5 | 417 | 281 | 27.04±8.76 | 31.67±11.09 | 41.29±18.44 |
| Synaptophysin | 382 | 250 | 34.59±6.83 | 30.17±6.27 | 35.24±12.31 |
| APP | 386 | 295 | 41.81±7.31 | 35.41±6.33 | 22.78±9.69 |

**Supplementary Table 3. Number of axons and tracks analyzed using Imaris software related to segmental analysis in Figure 2C, D, and E.**

| Axonal cargo | Nº moving tracks | Reversal frequency (nº/s) (mean±S.D.) | Pause frequency (nº/s) (mean±S.D.) | * Track Length (mean±S.D.) |
|---|---|---|---|---|
| TDP-43 | 266 | 1.44±0.31 | 0.67±0.31 | 13.37±4.43 |
| Rab5 | 281 | 1.13±0.43 | 1.10±0.34 | 9.89±3.15 |
| Synaptophysin | 250 | 1.18±0.25 | 1.02±0.32 | 12.96±3.75 |
| APP | 295 | 1.19±0.27 | 0.69±0.30 | 17.55±4.24 |

$$* \boldsymbol{TL} = \sum_{i=1}^{n-1} \sqrt{(x_{i+1} - x_i)^2 + (y_{i+1} - y_i)^2}$$

**Supplementary Table 4. Correlation analysis of segmental velocity *vs* segment duration.** The table shows specific rho Spearman's values for each protein, where segmental velocities significantly correlate with segment duration for both anterograde and retrograde. Z-scores were analyzed to evaluate significantly anterograde- or retrograde-driven correlations.

| Axonal cargo | Direction | Spearman ρ | C.I. | *p* value | *n of segments* | Anterograde vs Retrogade | |
|---|---|---|---|---|---|---|---|
| | | | | | | *z*-score | *p*-value |
| TDP-43 | Anterograde | 0.130 | 0.07961 to 0.1796 | <0.001 | 1574 | 0.1622 | 0.871 **(ns)** |
| | Retrograde | 0.124 | 0.07378 to 0.1742 | <0.001 | 1567 | | |
| Rab5 | Anterograde | 0.130 | 0.08114 to 0.1780 | <0.001 | 1680 | -1.061 | 0.288 **(ns)** |
| | Retrograde | 0.164 | 0.1206 to 0.2068 | <0.001 | 2075 | | |
| SYP | Anterograde | 0.290 | 0.2477 to 0.3319 | <0.001 | 1929 | 6.501 | 8E-11(***) |
| | Retrograde | 0.085 | 0.03798 to 0.1325 | <0.001 | 1796 | | |
| APP | Anterograde | 0.350 | 0.3078 to 0.3895 | <0.001 | 1881 | 3.154 | 0.00161 (**) |
| | Retrograde | 0.254 | 0.2085 to 0.2990 | <0.001 | 1742 | | |

### Supplementary Figure 2

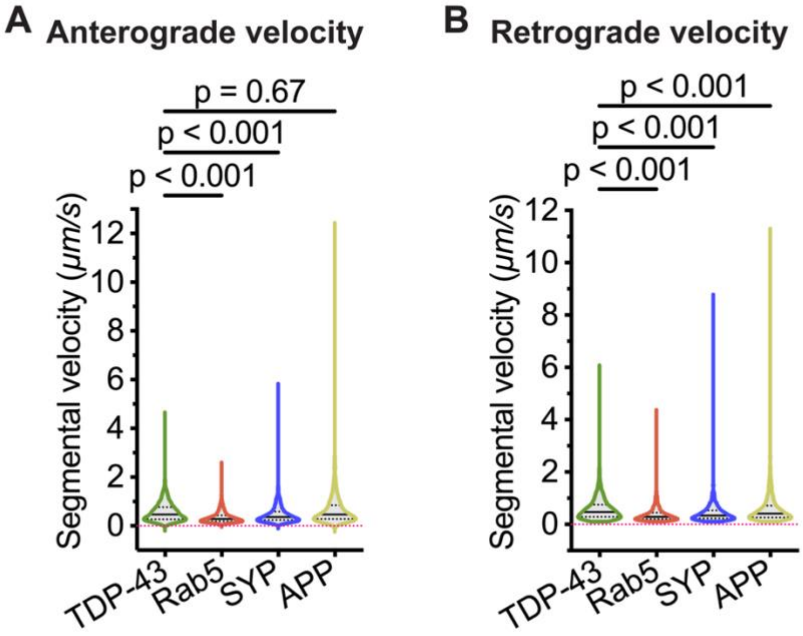


**Supplementary Figure 2 Segmental velocity correlation and distributions. (A)** Violin plots show overall distributions of anterograde segmental velocities among all the cargoes (*n* > 1500 segments – see table in **(B)** – were compared across the 4 proteins).

**(C)** Violin plots show overall distributions of retrograde segmental velocities among all the cargoes (*n* > 1500 segments – see table in **(A)** – were compared across the 4 proteins).

Data are shown as violin plots with median (middle bold line) and statistical analysis was performed with Kruskal–Wallis test followed by Dunn's multiple-comparisons post hoc test.

### Supplementary Figure 3

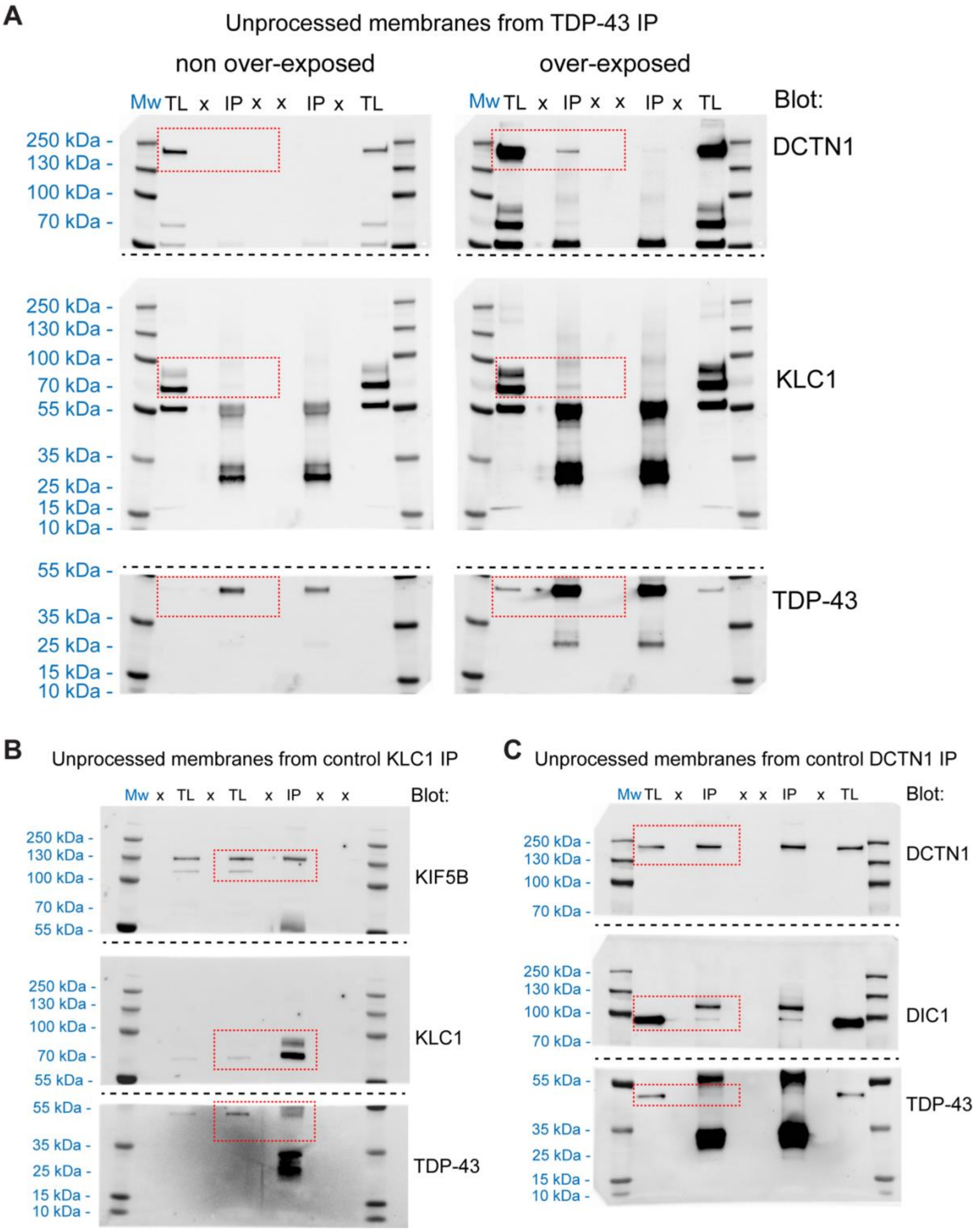

**Supplementary figure 3 Supporting unprocessed Western Blot images show Co-IP bands of IP-TDP43 with main anterograde and retrograde adaptors.**

**(A)** Unprocessed membranes of IP samples obtained from TDP-43 immunoprecipitations (IPs) prepared with cell lysates from neurons at DIV 40. TDP-43 blots were followed by KLC1, and DCTN1 to evaluate relative binding. The dashed red box highlights the area of the blot shown in **Figure 4B**. **(B)** Unprocessed membranes of IP samples obtained from KLC1 IP prepared as in (A). KLC1 blots were followed by TDP-43, and KIF5B blots to evaluate relative binding; KIF5B was used as a positive control of binding for KLC1. The dashed red box highlights the area of the blot shown in **Figure 4D**. **(C)** Unprocessed membranes of IP samples obtained from DCTN1 IP prepared as in **(A)**. DCTN1 blots were followed by TDP-43, and DIC1 blots to evaluate relative binding; DIC1 was used as a positive control of binding for DCTN1. The dashed red box highlights the area of the blot shown in **Figure 4F**.

For **(A)**, **(B)**, and **(C)**, the protein ladder was always imaged soon after the exposure of the bands of interest, and then the images merged for the proper marker-molecular weight correlation. The (x) indicate wells loaded with sample buffer only, while TL indicates the Total Lysate fraction (1% of IP=500μg).

### Supplementary Figure 4

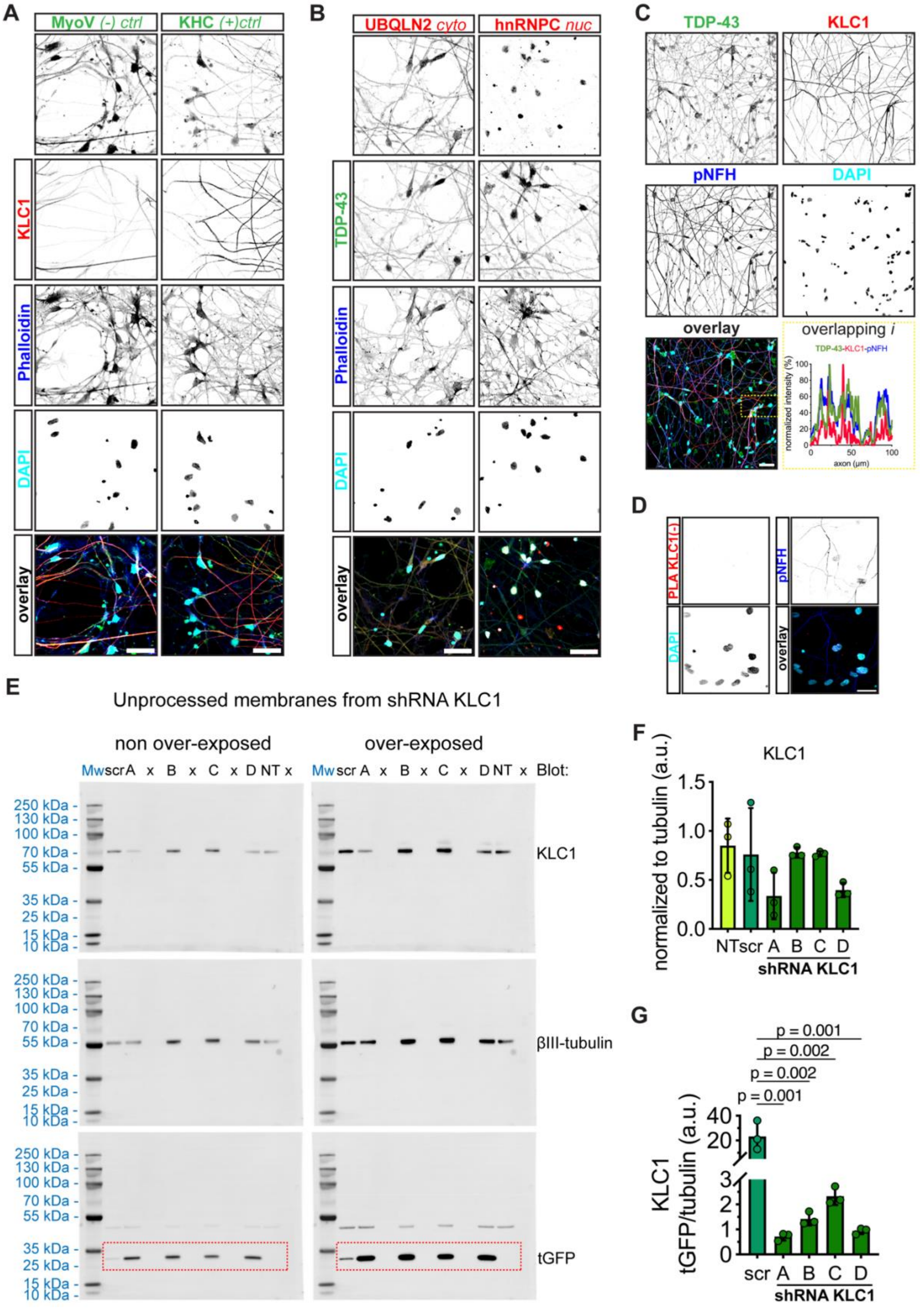

**Supplementary Figure 4 Overlapping signal of KLC1 and TDP-43 and effective downregulation of KLC1 expression in mature neurons. (A)** Representative micrographs of DIV 40 mature neurons stained for either MyoV or KHC (green) as negative and positive control as KLC1 interactors, respectively, then KLC1 (red), phalloidin (blue), and DAPI (cyan - scale bar = 50 μm). **(B)** Representative micrographs of DIV 40 mature neurons stained for either UBQLN2 or hnRNPC (red) as cytosolic and nuclear control as TDP-43 interactors, respectively, then TDP-43 (green), phalloidin (blue), and DAPI (cyan - scale bar = 50 μm). **(C)** Representative micrographs of DIV 40 mature neurons showing axonal overlapping of TDP-43 (green) and KLC1 (red) with pNFH (blue); DAPI shows nuclear stain (cyan - scale bar = 50 μm); normalized intensity graph (*lower right*) represents overlapping signal across TDP-43, KLC1, and pNFH, highlighted in the yellow dashed box region in the overlay image (*lower left*). **(D)** Representative micrographs of DIV 40 mature neurons showing negative control PLA for the KLC1 antibody (red), and the non-overlapping pNFH (blue); DAPI shows nuclear stain (cyan - scale bar = 50 μm). **(E)** Unprocessed membranes showing KLC1, βIII-tubulin, and turboGFP protein levels; lysates were obtained from DIV 40 mature neurons transduced with scramble_tGFP controls (scr), different shRNA_tGFP to downregulate KLC1 levels (A, B, C, and D), and non-transduced control samples (NT); blots on the left represent non over-exposed signals used for quantification, blots on the right show over-exposed protein signals to improve band visualization; red dashed boxes on tGFP blots highlight the latter, to do not confuse with upper stripped βIII-tubulin signals; 10 μg of proteins per each sample described above were loaded into gels. **(F)** Normalized quantification of protein levels for NT, scramble, and shRNA (A, B, C, and D) with βIII-tubulin loading control (*n*=3 biological replicate). **(G)** Normalized ratios of shRNA_tGFP efficacy in comparison with tGFP levels; each shRNA_tGFP KLC1 was compared with scramble_tGFP control (*n*=3 biological replicate).

Data show mean intensities of particles over 100 μm axonal distance (yellow dashed box) (**B**), and mean ± S.D. **(F, and G)**. Statistical comparison was performed using a One-way ANOVA followed by Dunnet's multiple comparisons test **(F -** *p*=ns**; G -** ***p*<0.01).

### Supplementary Figure 5

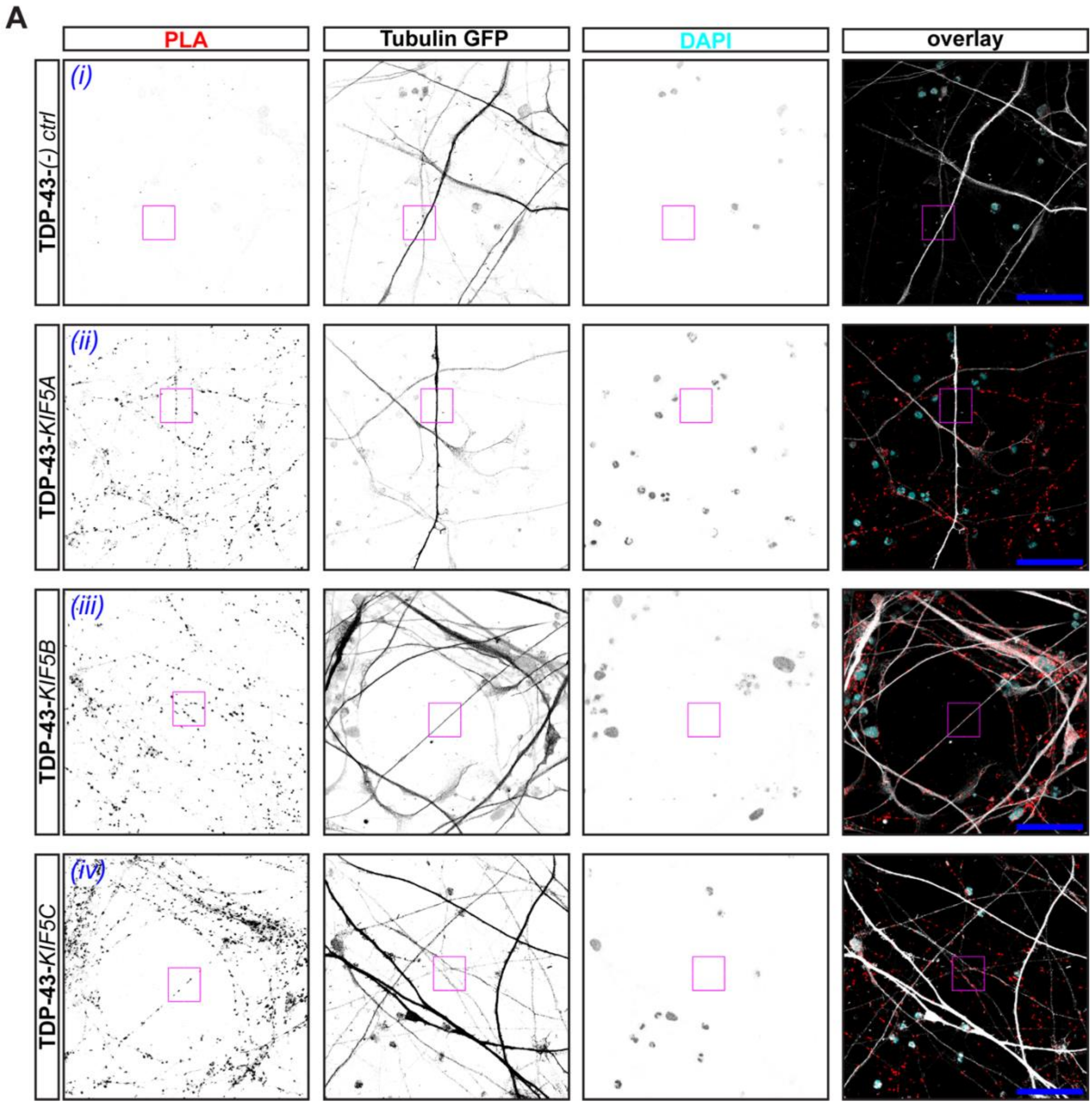


**Supplementary Figure 5 TDP-43 engages kinesin-1 family motors throughout axonal projections (A)** Representative micrographs of DIV 40 mature neurons unprocessed images showing negative control *(i)*, TDP-43-KIF5A *(ii)*, TDP-43-KIF5B *(iii)*, and TDP-43-KIF5C *(iv)* PLA staining. Micrographs show PLA dot signal (red), tubulin_GFP (white), and DAPI showing nuclear stain (cyan); dashed boxes highlight ROIs selected for zoom in micrograph representations in **Figure 5E** (scale bar = 50 μm).

### Supplementary Figure 6

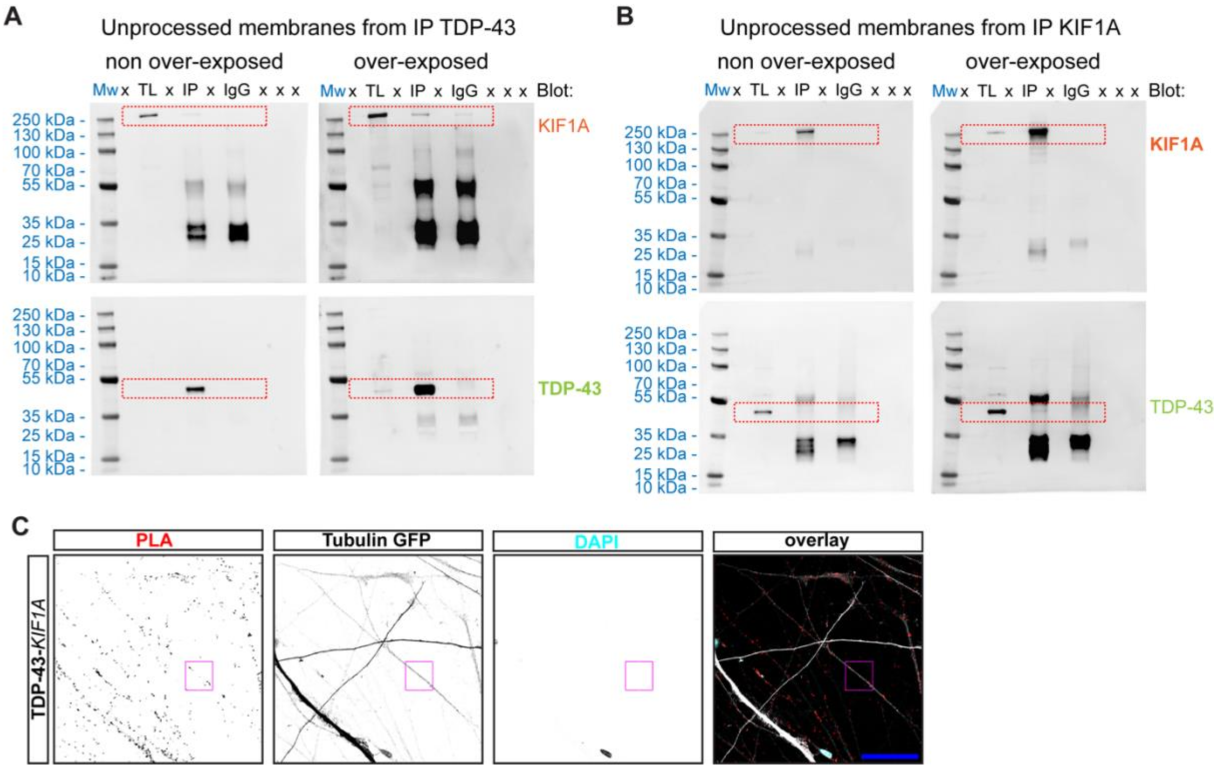


**Supplementary figure 6 TDP-43 associates with the synaptic vesicle motor KIF1A. (A)** Unprocessed membranes of IP samples obtained from TDP-43 immunoprecipitations (IPs) prepared with cell lysates from neurons at DIV 40. TDP-43 blots were followed by KIF1A to evaluate relative binding. The dashed red box highlights the area of the blot shown in **Figure 6B**, and the IgG isotype control for the TDP-43 antibody used for all the IP experiments in this study. **(B)** Unprocessed membranes of IP samples obtained from KIF1A IP prepared as in **(A)**. KLC1 blots were followed by TDP-43 blot to evaluate relative binding. The dashed red box highlights the area of the blot shown in **Figure 6C**, and the IgG isotype control for KIF1A antibody used for all the IP experiments in this study. For **(A)**, and **(B)**, the protein ladder was always imaged soon after the exposure of the bands of interest, and then the images merged for the proper marker-molecular weight correlation. The (x) indicate wells loaded with sample buffer only, while TL indicates the Total Lysate fraction (1% of IP=500µg and IgG=500µg). **(C)** Representative micrographs of DIV 40 mature neurons unprocessed images showing TDP-43-KIF1A interaction by the PLA dot signal (red), colocalizing with tubulin_GFP (white), and DAPI showing nuclear stain (cyan); dashed boxes highlight ROIs selected for zoom in micrograph representations in **Figure 6F** (scale bar = 50 µm).

### Supplementary movies legends

**Movie 1 – Transport in pNFH (+) axons of TDP-43_GFP transfected human neurons.**
Video of **TDP-43_GFP** particles recorded using a 489 nm laser at 4 fps, for a total of 240 frames: directionality of movement left (proximal to the cell body) to right (distal from the cell body). Scale bar = 10 µm.

**Movie 2 – Transport in pNFH (+) axons of GFP_Rab5 transfected human neurons.**
Video of **GFP_Rab5** particles recorded using a 489 nm laser at 4 fps, for a total of 240 frames: directionality of movement left (proximal to the cell body) to right (distal from the cell body). Scale bar = 10 µm.

**Movie 3 – Transport in pNFH (+) axons of Synaptophysin_GFP transfected human neurons.**
Video of **Synaptophysin_GFP** particles recorded using a 489 nm laser at 4 fps, for a total of 240 frames: directionality of movement left (proximal to the cell body) to right (distal from the cell body). Scale bar = 10 µm.

**Movie 4 – Transport in pNFH (+) axon s of APP_GFP transfected human neurons.**
Video of **APP_GFP** particles recorded using a 489 nm laser at 4 fps, for a total of 240 frames: directionality of movement left (proximal to the cell body) to right (distal from the cell body). Scale bar = 10 µm.
